# Identifying and prioritizing potential human-infecting viruses from their genome sequences

**DOI:** 10.1101/2020.11.12.379917

**Authors:** Nardus Mollentze, Simon A. Babayan, Daniel G. Streicker

## Abstract

Determining which animal viruses may be capable of infecting humans is currently intractable at the time of their discovery, precluding prioritization of high-risk viruses for early investigation and outbreak preparedness. Given the increasing use of genomics in virus discovery and the otherwise sparse knowledge of the biology of newly-discovered viruses, we developed machine learning models that identify candidate zoonoses solely using signatures of host range encoded in viral genomes. Within a dataset of 861 viral species with known zoonotic status, our approach outperformed models based on the phylogenetic relatedness of viruses to known human-infecting viruses (AUC = 0.773), distinguishing high-risk viruses within families that contain a minority of human-infecting species and identifying putatively undetected or so far unrealized zoonoses. Analyses of the underpinnings of model predictions suggested the existence of generalisable features of viral genomes that are independent of virus taxonomic relationships and that may preadapt viruses to infect humans. Our model reduced a second set of 645 animal-associated viruses that were excluded from training to 272 high and 41 very high-risk candidate zoonoses and showed significantly elevated predicted zoonotic risk in viruses from non-human primates, but not other mammalian or avian host groups. A second application showed that our models could have identified SARS-CoV-2 as a relatively high-risk coronavirus strain and that this prediction required no prior knowledge of zoonotic SARS-related coronaviruses. Genome-based zoonotic risk assessment provides a rapid, low-cost approach to enable evidence-driven virus surveillance and increases the feasibility of downstream biological and ecological characterisation of viruses.

## Introduction

Most emerging infectious diseases of humans are caused by viruses that originate from other animal species. Identifying these zoonotic threats prior to emergence is a major challenge since only a small minority of the estimated 1.67 million animal viruses may infect humans [1–3]. Existing models of human infection risk rely on viral phenotypic information that is unknown for newly discovered viruses (e.g., the diversity of species a virus can infect) or that vary insufficiently to discriminate risk at the virus species or strain level (e.g., replication in the cytoplasm), limiting their predictive value before the virus in question has been characterised [4–6]. Since most viruses are now discovered using untargeted genomic sequencing, often involving many simultaneous discoveries with limited phenotypic data, an ideal approach would quantify the relative risk of human infectivity upon relevant exposure from sequence data alone. By identifying high risk viruses warranting further investigation, such predictions could alleviate the growing imbalance between the rapid pace of virus discovery and lower throughput field and laboratory research needed to comprehensively evaluate risk.

Current models can identify well-characterised human-infecting viruses from genomic sequences [7,8]. However, by training algorithms on very closely related viruses (i.e., strains of the same species) and potentially omitting secondary characteristics of viral genomes linked to infection capability, such models are less likely to find signals of zoonotic status that generalise across viruses. Consequently, predictions may be highly sensitive to substantial biases in current knowledge of viral diversity [3,9].

Empirical and theoretical evidence suggests that generalisable signals of human infectivity might exist within viral genomes. Viruses associated with broad taxonomic groups of animal reservoirs (e.g., primates versus rodents) can be distinguished using aspects of their genome composition, including dinucleotide, codon, and amino acid biases [10]. Whether such measures of viral genome composition are specific enough to distinguish host range at the species level remains unclear, but their specificity might arise through several commonly hypothesised mechanisms. First, aspects of antiviral immunity that target nucleotide motifs in viral genomes might select for common mutations in diverse human-associated viruses [11,12]. For example, the depletion of CpG dinucleotides in vertebrate-infecting RNA virus genomes may have arisen to evade zinc-finger antiviral protein (ZAP), an interferon-stimulated gene (ISG) that initiates the degradation of CpG-rich RNA molecules [12]. While ZAP occurs widely among vertebrates, increasingly recognized lineage-specificity in vertebrate antiviral defences opens the possibility that analogous, undescribed nucleic-acid targeting defences might be human (or primate) specific [13]. Second, the frequencies of specific codons in virus genomes often resemble those of their reservoir hosts, possibly owing to increased efficiency and/or accuracy of mRNA translation [14]. By driving genome compositional similarity to human-adapted viruses or to the human genome, such processes may preadapt viruses for human infection [15,16]. Finally, even in the absence of mechanisms that assert common selective pressures on divergent viral genomes, the phylogenetic relatedness of viruses could allow prediction of the potential for human infectivity since closely related viruses are generally assumed to share common phenotypes and host range. However, despite being a common rule of thumb for virus risk assessment, to our knowledge, whether evolutionary proximity to viruses with known human infection ability predicts zoonotic status remains untested.

We aimed to develop machine learning models which use features engineered from viral and human genome sequences to predict the probability that any animal-infecting virus will infect humans given biologically relevant exposure (here, zoonotic potential). Using a large dataset of viruses which had previously been assessed for human infection ability based on published reports, we first build models which assess zoonotic potential based on virus taxonomy and/or phylogenetic relatedness to known human-infecting viruses and contrast these models to alternatives based on hypothesized selective pressures on viral genome composition which favour human infectivity. We then apply the best-performing model to explore patterns in the predicted zoonotic potential of additional virus genomes sampled from a range of species.

## Results

We collected a single representative genome sequence from 861 RNA and DNA virus species spanning 36 viral families that contain animal-infecting species (S1 Fig). We labelled each virus as being capable of infecting humans or not using published reports as ground truth, and trained models to classify viruses accordingly. These classifications of human infectivity were obtained by merging three previously published datasets which reported data at the virus species level and therefore did not consider potential for variation in host range within virus species [5,9,17]. Importantly, given diagnostic limitations and the likelihood that not all viruses capable of human infection have had opportunities to emerge and be detected, viruses not reported to infect humans may represent unrealized, undocumented, or genuinely non-zoonotic species. Identifying potential or undocumented zoonoses within our data was an *a priori* goal of our analysis.

We first evaluated whether phylogenetic proximity to human-infecting viruses predictably elevates zoonotic potential. Gradient boosted machine (GBM) classifiers trained on virus taxonomy or the frequency of human-infecting viruses among close relatives identified by sequence similarity searches (“phylogenetic neighbourhood”, defined using nucleotide BLAST [10]) outperformed chance (median area under the receiver-operating characteristic curve [AUC_m_] = 0.604 and 0.558, respectively), but were no better than manually ranking novel viruses by the proportion of human-infecting viruses in each family (“taxonomy-based heuristic”, AUC_m_ = 0.596, Fig 1A). This indicates that relatedness-based models were not only unable to identify novel zoonoses which are not close relatives of known human-infecting viruses, but were also largely unable to accurately distinguish risk among closely related viruses (S2 Fig). Moreover, the performance of these models depended on the data available for model training, sometimes performing worse than chance, making them highly sensitive to current knowledge of viral diversity.

**Fig 1.**
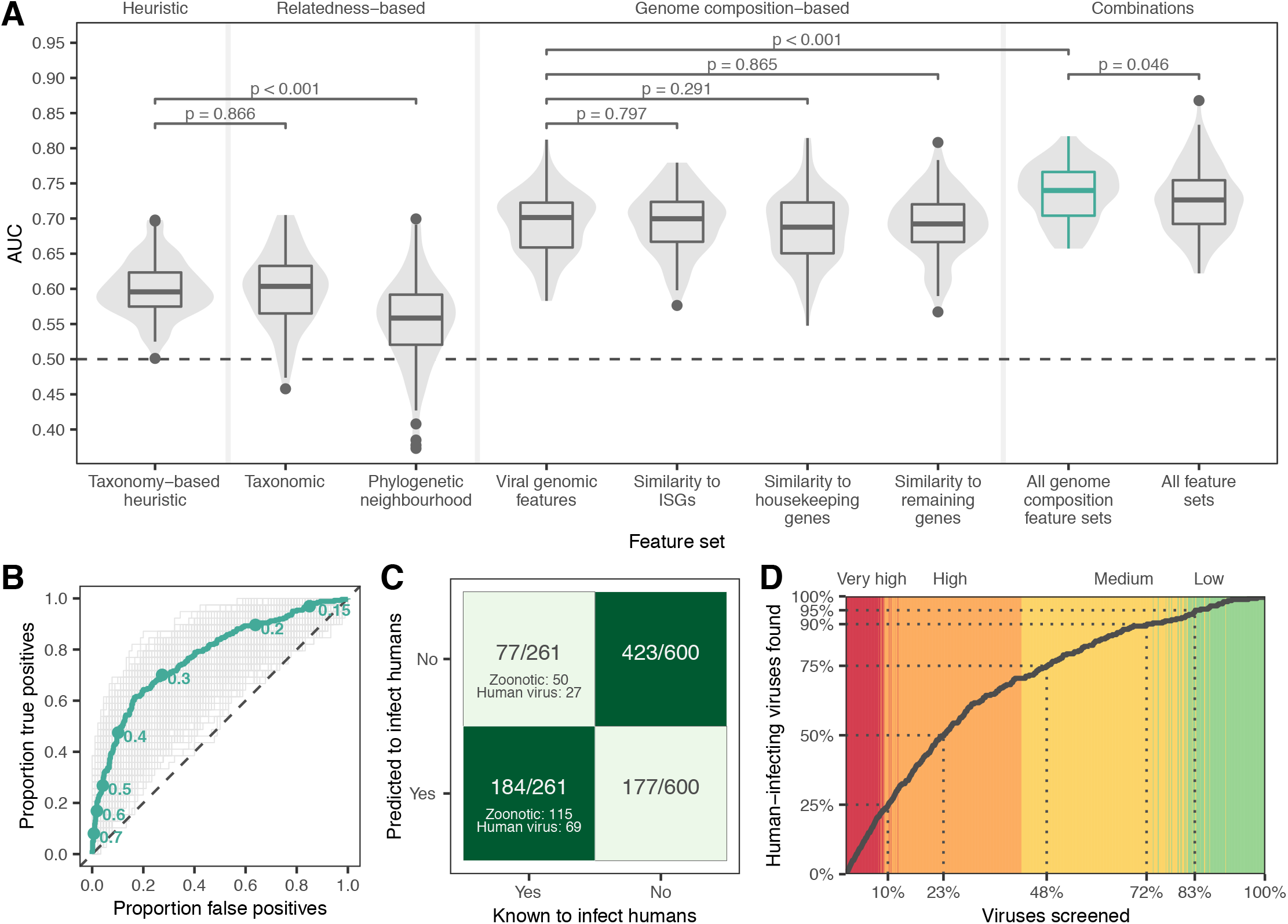
Machine learning prediction of human infectivity from viral genomes. (A) Violins and boxplots show the distribution of AUC scores across 100 replicate test sets. (B) Receiver-operating characteristic curves showing the performance of the model trained on all genome composition feature sets across 1000 iterations (grey) and performance of the bagged model derived from the top 10% of iterations (green). Points indicate discrete probability cut-offs for categorizing viruses as human-infecting. (C – D) show binary predictions and discrete zoonotic potential categories from the bagged model, using the cut-off which balanced sensitivity and specificity (0.293). (C) Heatmap showing the proportion of predicted viruses in each category. (D) Cumulative discovery of human-infecting species when viruses are prioritised for downstream confirmation in the order suggested by the bagged model. Dotted lines highlight the proportion of all viruses in the training and evaluation data which need to be screened to detect a given proportion of known human-infecting viruses. Background colour highlights the assigned zoonotic potential categories of individual viruses encountered (red: very high, orange: high, yellow: medium, green: low).

We next quantified the performance of GBMs trained on genome composition (i.e., codon usage biases, amino acid biases, and dinucleotide biases), calculated either directly from viral genomes (“viral genomic features”) or based on the similarity of viral genome composition to that of three distinct sets of human gene transcripts (“human similarity features”): interferon-stimulated genes (ISGs), housekeeping genes, and all other genes. We hypothesized that if viruses need to adapt to either evade innate immune surveillance for foreign nucleic acids or to optimise gene expression in humans, they should resemble ISGs since both tend to be expressed concomitantly in virus-infected cells. We selected two additional sets comprising non-ISG housekeeping genes and all remaining genes to explore whether signals were specific to ISGs. GBMs trained using genome composition feature sets performed similarly when tested separately (AUC_m_ = 0.688–0.701) and consistently outperformed models based on relatedness alone (both the taxonomy-based heuristic and machine learning models trained on virus taxonomy or phylogenetic neighbourhood, Fig 1A). Combining all four genome composition feature sets further improved, and reduced variance in, performance (AUC_m_ = 0.740, Fig 1A), suggesting that measures of similarity to human transcripts contained information unavailable from viral genomic features alone. In contrast, adding relatedness features to this combined model reduced accuracy (AUC_m_ = 0.726) and increased variance (Fig 1A). Averaging output probabilities over the best 100 out of 1000 iterations of training on random test/train splits of the data (a process akin to bagging, using ranking performance on non-target viruses to select high performing models) further improved the combined genome feature-based model (AUC = 0.773, Fig 1B).

To estimate model sensitivity and specificity, we converted the mean of predicted probabilities of human infection from the bagged model into binary classifications (i.e., human infecting or not), predicting viruses with predicted probabilities > 0.293 as human-infecting. This cut-off balanced sensitivity and specificity (both 0.705, Fig 1C), though in principle, higher or lower cut-offs could be selected to prioritize reduction of false positives or false negatives, respectively (Fig 1B). These binary predictions correctly identified 71.9% of viruses which predominately or exclusively infect humans and 69.7% of zoonotic viruses as human infecting, though performance varied among viral families (Fig 1C, S3 Fig). Since binary classifications ignore both the variability between iterations and the rank of viruses relative to each other, we further converted predicted probabilities of zoonotic potential into four zoonotic potential categories, describing the overlap of confidence intervals with the 0.293 cut-off from above (low: entire 95% confidence interval [CI] of predicted probability ≤ cut-off; medium: mean prediction ≤ cut-off, but CI crosses it; high: mean prediction > cut-off, but CI crosses it; very high: entire CI > cut-off). Under this scheme, the majority (92%) of known human-infecting viruses were predicted to have either medium (21.5%), high (47.1%) or very high (23.4%) zoonotic potential, while only 8% (N=21) had low zoonotic potential (S4 Fig, S1 Table). Fourteen viruses not currently considered to infect humans by our criteria were predicted to have very high zoonotic potential (S5 Fig), although at least three of these (*Aura virus*, *Ndumu virus* and *Uganda S virus*) have serological evidence of human infection [5,17], suggesting they may be valid zoonoses rather than model misclassifications. Across the full dataset, 77.2% of viruses predicted to have very high zoonotic potential were known to infect humans (S1 Table). Consequently, studies aimed at confirming human infectivity (e.g., by attempting to infect human-derived cell lines or by serological testing of humans in high-risk populations) while screening viruses in the order suggested by our ranking would have found 23.4% of all known human-infecting viruses in this dataset after screening just the very high zoonotic potential viruses (9.2% of all viruses). More generally, 50% of known human-infecting viruses would have been found after screening the top-ranked 23.3% of viruses, and 75% after screening the top 48% of viruses (Fig 1D). In contrast, if relying only on relatedness to known zoonoses, confirming the first 50% of currently-known zoonoses would have required screening either 40.2% (taxonomy-based model) or 41.5% (phylogenetic neighbourhood-based model) of viruses; a 1.7 to 1.8-fold increase in effort compared to our best model (S6 Fig).

Since genome composition features partly track viral evolutionary history [10], it is conceivable that our models made predictions by reconstructing taxonomy more accurately than the phylogenetic neighbourhood estimator or in more detail than available to the taxonomy-based model. We therefore compared dendrograms that clustered viruses by either taxonomy, raw genomic features, or the relative influence of each genomic feature on the model prediction for each virus (here estimated using the SHAP algorithm of [18], and hence termed SHAP values in the rest of this manuscript). As high levels of similarity in SHAP values between viruses would indicate that they share similar predictions for the same reasons, we therefore analysed to what extent such similar uses of the same genomic features followed known taxonomic patterns. While dendrograms using raw feature values closely correlated with virus taxonomy for both human-infecting and other viruses (Baker’s [19] γ = 0.617 and 0.492 respectively, p < 0.001), dendrograms of SHAP similarity had 10.28- and 2.07-fold reduced correlations with virus taxonomy (γ = 0.060 and 0.238, although this was still more correlated than expected by chance, p ≤ 0.008; S7 Fig). Among human-infecting viruses, correlations between SHAP similarity-based clustering and virus taxonomy weakened at deeper taxonomic levels, even though the input genomic features provided sufficient information to partially reconstruct virus taxonomy at the realm, kingdom and phylum levels (S8 Fig). These results indicate that more taxonomic information was available than was utilised by the trained model to predict human infection ability. Interestingly, dendrograms of SHAP similarity showed that even viruses with different genome types – indicating ancient evolutionary divergence or separate origins – clustered together (Fig 2A). Alongside earlier observations on classifier performance (Fig 1A), this suggests that the genome composition-based model outperformed relatedness-based approaches because it found common viral genome features that increase the capacity for human infection across diverse viruses.

**Fig 2.**
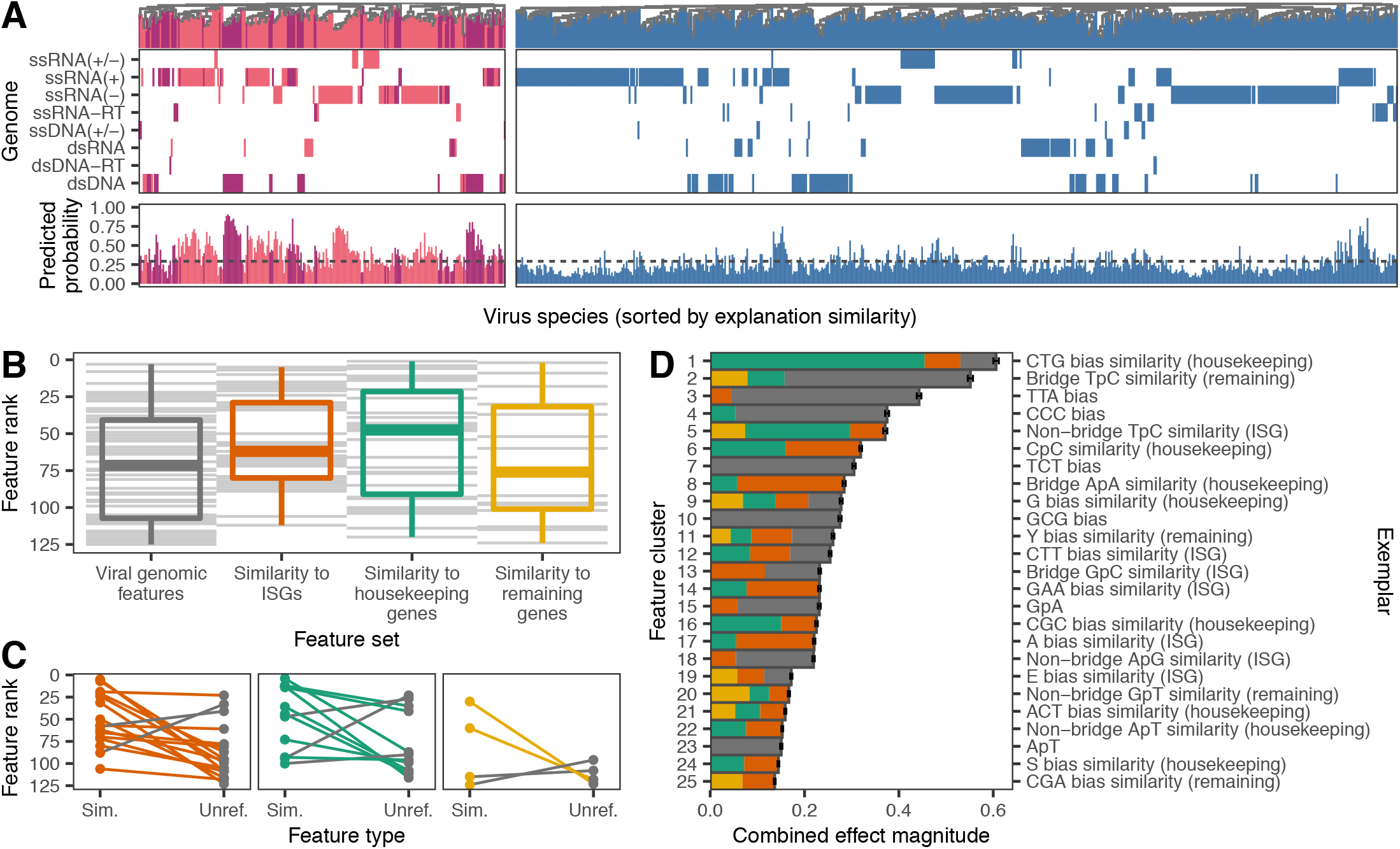
Genomic determinants of human-infecting viruses. (A) SHAP value clustering of viruses known to infect humans (primarily human-associated, dark purple, and zoonotic, pink) and those with no known history of human infection (blue) shows that similar features predicted human infection across viruses with different genome types (rows). A second set of panels shows the predicted probability of infecting humans for each virus, with the dashed line indicating the cut-off which balances sensitivity and specificity. (B) Relative importance of individual features in shaping predictions, determined by ranking features by the mean of absolute SHAP values across all viruses. Grey lines represent individual features; boxplots show the median, 25^th^/75^th^ percentiles and range of ranks for each feature set. (C) Difference in ranks of features when both unreferenced (“Unref.”) and similarity to human genomes (“Sim.”) forms were retained in the final model. Lines are coloured according to the highest ranked representation in each pairwise comparison; colours as in B. (D) Composition of the top 25 most important clusters of correlated features shaping predictions. Discrete clusters of correlated features were identified by affinity propagation clustering. Clusters are shown ranked by the combined effect magnitude of constituent features, defined as the sum of mean absolute SHAP values for all features in the cluster, and the exemplar feature of each cluster is provided on the right axis. Bars represent means (± SEM) across 1000 iterations and are shaded by the proportion of the cluster from each feature set; colours as in B.

Although our analysis was not designed to conclusively identify biological mechanisms underlying genomic predictors of human infection, we nevertheless were able to explore emergent patterns relating to how specific genome composition features and groups of features relate to human infectivity. We first compared the relative influence of features from different genome composition categories (i.e., genomic features versus the three sets of human similarity features). Representatives of all genome composition categories were retained in the final model, though we found some evidence that compositional similarity to human housekeeping genes and ISGs influenced predictions more strongly than unreferenced viral genomic features (Fig 2B–C; S1 Text). We next explored the influence of individual features on model predictions in more detail. Unsurprisingly, given that GBMs are designed to make predictions from large numbers of weakly informative features [20], no single feature stood out as the driving force and many features formed correlated clusters (Fig 2D & S9 Fig). More interestingly, many features had complex, non-linear relationships with human infection (S10 Fig), such that increased similarity to human gene transcripts did not always increase the likelihood of infecting humans (S1 Text). We speculate this might reflect trade-offs between different features within viral genomes or context dependencies whereby both mimicry of human transcripts (e.g., for improved translation efficiency) or divergence from human transcripts (e.g., for evasion of nucleotide motif-targeting defences) may occur for different features (S1 Text).

Finally, we carried out two case studies to illustrate the utility of our prediction framework. First, we used the combined genome feature-based model to rank 758 virus species which were not present in our training data. We included all species in the most recent ICTV taxonomy release (#35, 24 April 2020) belonging to animal-infecting virus families and which were originally discovered or sequenced from mammals (including humans), birds, two insect orders containing common virus vectors (Diptera and Ixodida), or where the sampled host was not reported. This dataset contained representatives from 38 viral families, including 2 (*Anelloviridae* and *Genomoviridae*) which were not present in data used to train our model. In total, 70.8% of viruses sampled from humans were correctly identified as having either very high (N=36) or high zoonotic potential (N=44; Fig 3A). The remaining human-associated viruses were primarily classified as medium zoonotic potential (N=30), with 3 species predicted to have low zoonotic potential (*Mammalian orthoreovirus* and *Human associated gemykibivirus 2* and *3*; Fig 3A). Within the viral families never previously seen by our model, the majority of human-associated anelloviruses (39/45, 86.6%) were correctly identified as having either very high or high zoonotic potential, consistent with the conclusion that viral genomic features which enhance human infectivity can generalise across viral families. In contrast, all 6 human-associated genomoviruses were classified as either medium or low zoonotic potential. The lower performance on genomoviruses may reflect the unusual genomic structure of this family (circular, single stranded DNA) which was poorly represented in training (only 2 representatives from the *Circoviridae* family; S1 Fig) and may impose different selective forces. Further, the small genome sizes of genomoviruses (2.2– 2.4kb) may complicate calculation of genomic features due to the low number of nucleotides, dinucleotides and codons available (cf. S3 Fig). Among the 645 viruses with unknown human infectivity that were sequenced from non-human animal or potential vector samples, 45.0% were predicted to have either very high (N=41) or high zoonotic potential (N=272; S11 Fig & S1 Table). The very high zoonotic potential category was dominated by *Papillomaviridae* (34.1%) and *Peribunyaviridae* (19.5%).

**Fig 3.**
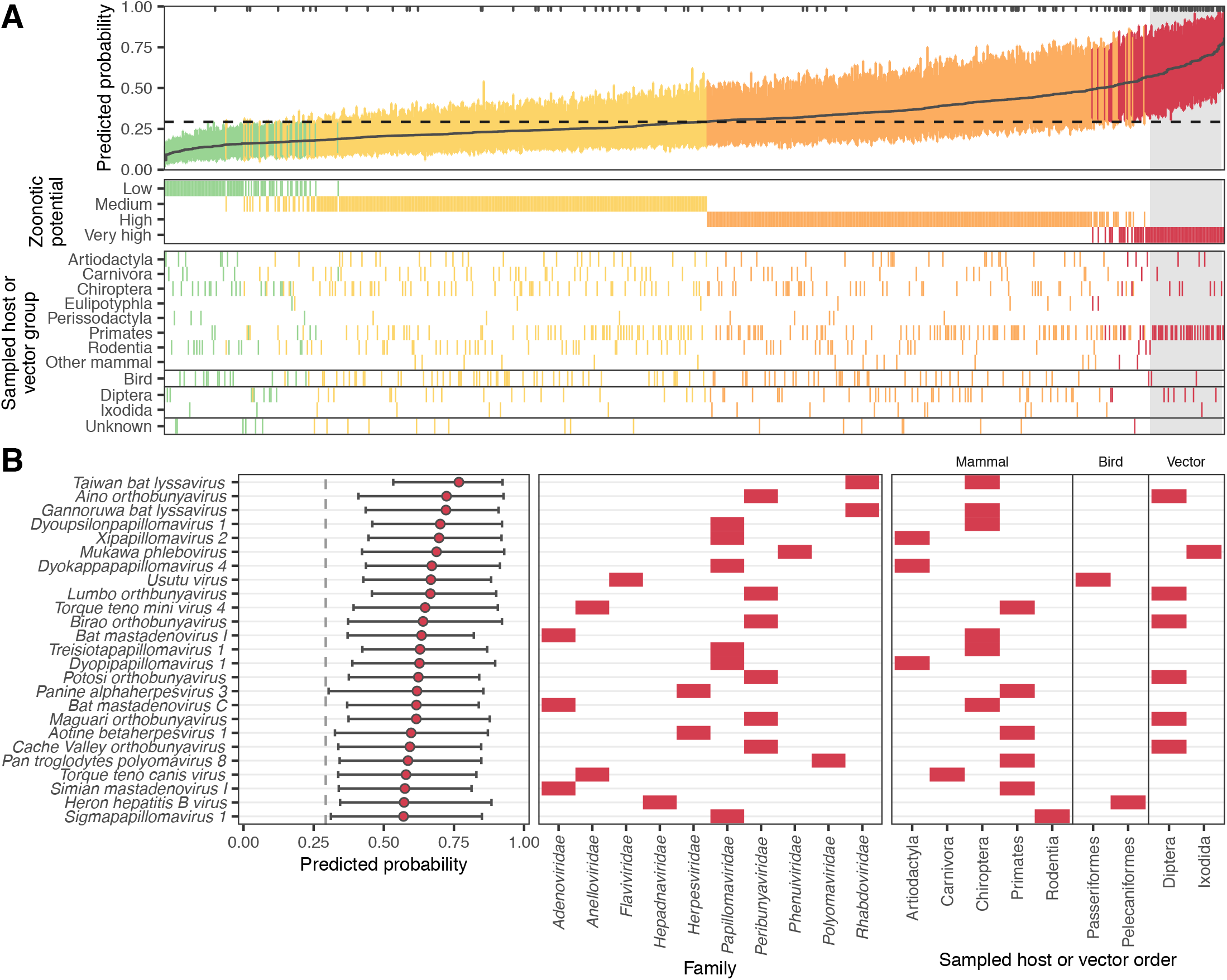
Probability of human infection predicted from holdout viral genomes. (A) Predicted probability of human infection for 758 virus species which were not in the training data. Colours show the assigned zoonotic potential categories, with an additional panel showing the host or vector group each virus genome was sampled from. Tick marks along the top edge of the first panel show the location of virus genomes sampled from humans, while a dashed line shows the cut-off which balanced sensitivity and specificity in the training data. The top 25 viruses which were not sampled from humans (contained within the grey box) are illustrated in more detail in (B). Bars show the 95% interquantile range of predicted probabilities across the best-performing 10% of iterations (based on the training data), while a solid line (A) or circles (B) show the mean predicted probability from these iterations.

We next used a beta regression model to explore how predictions of zoonotic potential varied among host and viral groups. As expected given the performance on our training and evaluation data (Fig 1), the 113 virus species that were sequenced from human samples scored consistently higher than those detected in other hosts (p < 0.001; figs. 3A & 4D). Although viruses from putatively high risk host groups including bats, rodents and artiodactyls formed a large fraction of our holdout data (with viruses from bats outnumbering even those from humans, S11 Fig), they did not have elevated predicted probabilities of being zoonotic (Fig 4C), and no differences were detected at higher host taxonomic levels (Fig 4A – B). This highlights a potential disparity between current sampling efforts for virus discovery/reporting and the distribution of zoonotic risk. In contrast, viruses linked to primates had higher predicted probabilities of infecting humans, even after accounting for human-associated viruses and the effects of virus family (Figs 3 – 4 & S11 Fig). That genome composition-based models predicted elevated zoonotic potential in non-human primate-associated viruses despite receiving no information on sampled host further supports host-mediated selective processes as a biological basis for our model’s predictions. In addition to relatively rare and small host effects, we observed more pervasive positive and negative effects of virus family on predicted zoonotic status (Fig 4E). Taken together, our results are consistent with the expectation that the relatively close phylogenetic proximity of non-human primates may facilitate virus sharing with humans and suggest that this may in part reflect common selective pressures on viral genome composition in both humans and non-human primates. However, broad differences among other animal groups appear to have less influence on zoonotic potential than virus characteristics [9].

**Fig 4.**
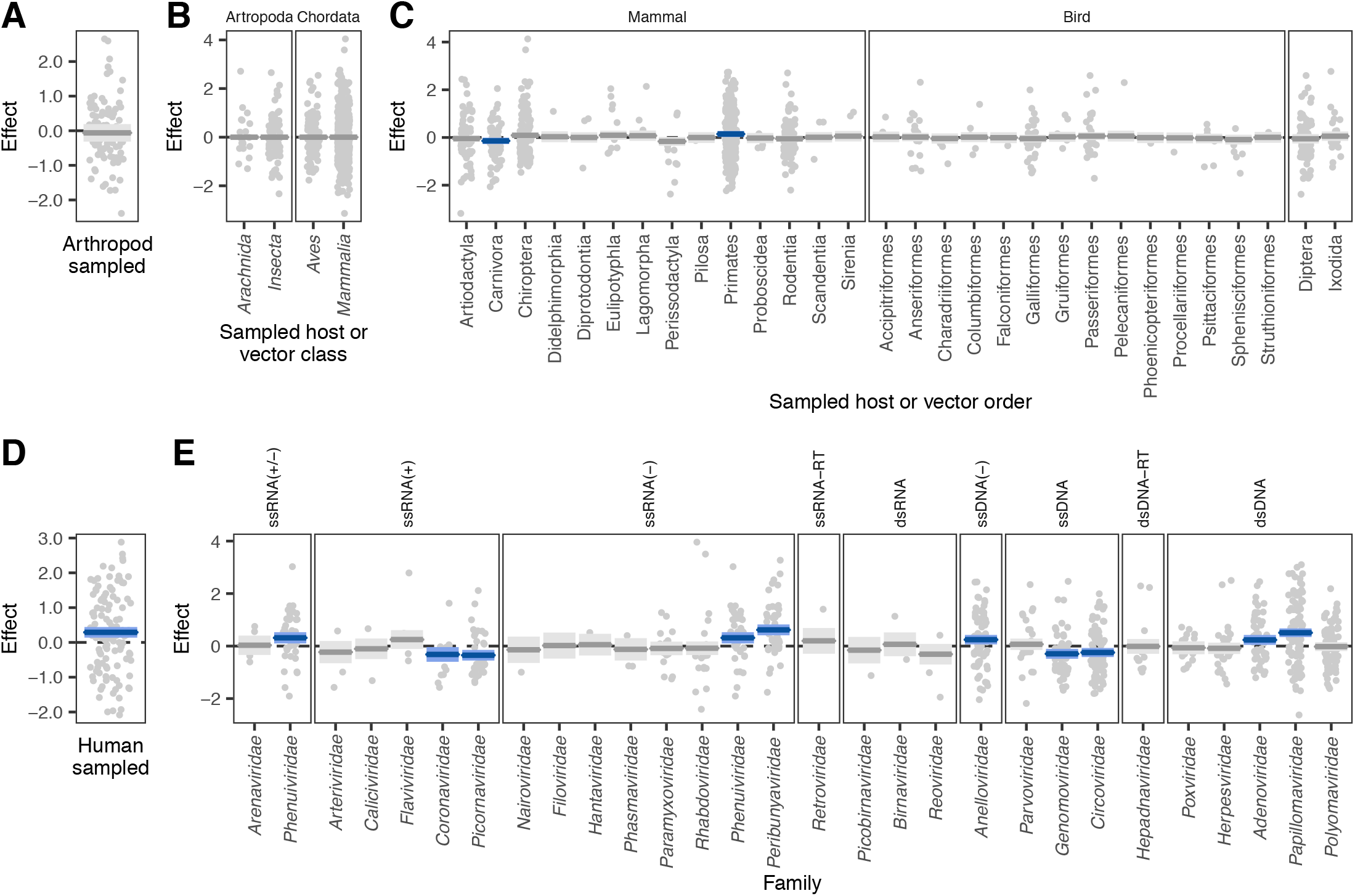
Factors correlated with the probability of human infection predicted from holdout viral genomes. Partial effects plots are shown for a beta regression model attempting to explain the mean probability assigned by the bagged model to all viruses in Fig 3A, accounting for whether or not the genome predicted was sequenced from arthropods (as opposed to chordates, A), random effects for the taxonomic class and order of sampled hosts (B – C), whether the sequence derived from a human sample (D), and a random effect for the virus family represented (E). Points indicate partial residuals, while lines and shaded areas respectively show the maximum likelihood and 95% confidence interval of partial effects. Confidence intervals which do not include zero are highlighted in blue.

Our second case study used coronaviruses to explore the ability of our combined genome feature-based model to distinguish different virus species within the same family and different genomes within a single virus species. Specifically, we predicted the zoonotic potential of all currently recognized coronavirus species, along with 62 human and animal-derived genomes of *Severe acute respiratory syndrome-related coronavirus* (SARS coronavirus), a species which includes many well-known viruses, including SARS-CoV, SARS-CoV-2, and related sarbecoviruses. All known human-infecting coronaviruses were classified as either medium or high zoonotic potential (Fig 5A). We also identified two additional animal-associated coronaviruses – *Alphacoronavirus 1* and the recently described *Sorex araneus coronavirus T14* – as being at least as, or more likely to be capable of infecting humans than known, high ranking, human-infecting coronaviruses; these should be considered high priority for further research. While this manuscript was in revision, a recombinant *Alphacoronavirus 1* was detected in nasopharyngeal swabs from pneumonia patients, further strengthening the case that this species may be zoonotic [21]. We further observed variation in predicted zoonotic potential within coronavirus genera which was consistent with our current understanding of these viruses. *Alphacoronavirus* and *Betacoronavirus* (the genera which contain known human-infecting species) also contained non-zoonotic species that were correctly predicted to have low zoonotic potential, while the majority of delta- and gammacoronaviruses received relatively low predictions (Fig 5A). These findings further illustrate the capacity of our models to discriminate risk below the virus family or genus levels.

**Fig 5.**
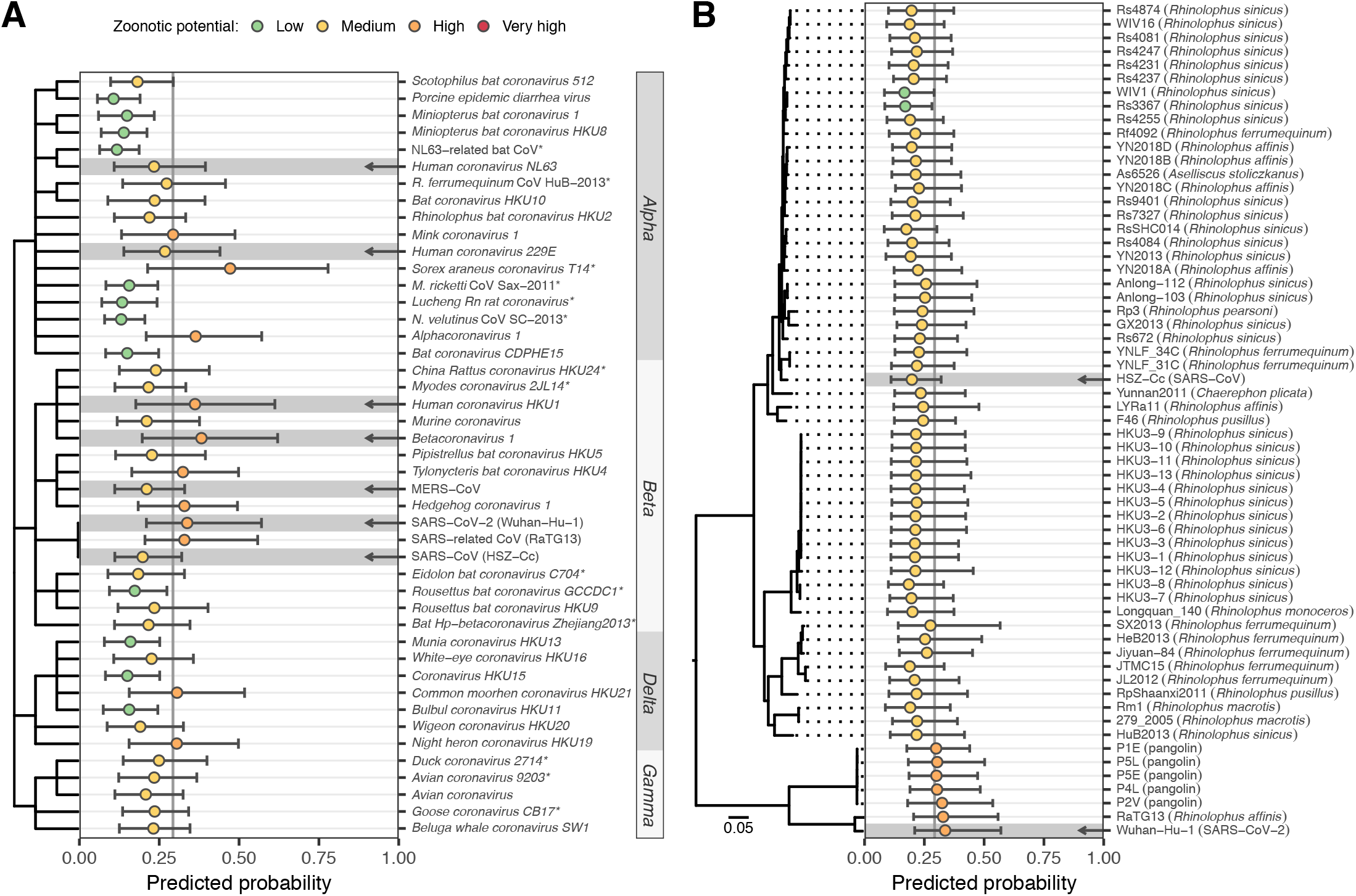
Probability of human infection predicted from Coronavirus genomes. (A) Predictions for currently recognized *Coronaviridae* species and for 3 variants of *Severe Acute Respiratory Syndrome-related coronavirus*: SARS-CoV (isolate HSZ-Cc, sampled early in the 2003 pandemic), SARS-CoV-2 (isolate Wuhan-Hu-1, sampled early in the current pandemic), and the closely related RaTG13 (sampled from *Rhinolophus affinis* in 2013). A dendrogram illustrates taxonomic relationships, with abbreviated genus names annotated on the right. Arrows highlight known human-infecting species. Asterisks indicate species absent from the training data, also present in Fig 3A. (B) Predictions for different representatives of *Severe acute respiratory syndrome-related coronavirus*. The isolation source of animal-associated genomes is indicated in parentheses. A maximum likelihood phylogeny illustrates relationships and was created as described in [6]. The outgroup, BtKy72 (sampled in Kenya in 2007), is not shown. MERS-CoV = *Middle East respiratory syndrome-related coronavirus*; *M. ricketti* CoV Sax-2011 = *Myotis ricketti alphacoronavirus Sax-2011*; NL63-related bat CoV = *NL63-related bat coronavirus strain BtKYNL63-9b*; *N. velutinus* CoV SC-2013 = *Nyctalus velutinus alphacoronavirus SC-2013*; *R. ferrumequinum* CoV HuB-2013 = *Rhinolophus ferrumequinum alphacoronavirus HuB-2013*. In both panels, bars show the 95% interquantile range of predicted probabilities across the best-performing 10% of iterations excluding the species being predicted, while circles show the mean predicted probability from these iterations.

Within SARS coronavirus, most genomes (85.5%) were classified as having medium zoonotic potential, including the causal agent of the 2003 outbreak (Fig 5B). Interestingly however, SARS-CoV-2 (the causative agent of the current COVID-19 pandemic), the closely related RaTG13 from a rhinolophid bat, and all 5 closely-related pangolin-associated isolates tested were predicted to have high zoonotic potential (although confidence intervals between all sarbecoviruses tested overlapped, Fig 5B). Importantly, these predictions were made using iterations of our model which excluded the 2003 SARS-CoV genome or any other sarbecovirus from training. This finding, together with our observation that relatively few other animal-infecting, allegedly non-zoonotic coronaviruses had similarly high scores, suggests that the elevated risk of SARS-CoV-2 and closely-related genomes discovered in animals could have been anticipated via sequencing-based surveillance and might have led to actionable research or surveillance prior to the zoonotic emergence of any sarbecovirus (Fig 5).

## Discussion

In an age of rapid, genomic-based virus discovery, rational prioritization of research and surveillance activities has been an unresolved challenge. While approaches to prioritise known, relatively well-characterised viruses based on a range of common risk factors have been developed [4–6], the large number of viruses still being discovered presents a bottleneck for the very characterisation needed to apply such prioritisation schemes, necessitating use of expert opinion or surrogate data from related species [6]. Our findings show that the zoonotic potential of viruses can be inferred to a surprisingly large extent from their genome sequence, outperforming current alternatives. Indeed, our results suggest that routine proxies of zoonotic risk which can be applied to poorly characterised viruses including virus taxonomy and relative phylogenetic proximity to human-infecting species [5,9,22] have limited discriminatory power. This has far-reaching implications for how risk is perceived – while it is intuitive to assume that novel viruses which are closely related to known human-infecting viruses are a threat, to our knowledge this assumption had never been tested. Worryingly, with some training datasets such relatedness-based models performed worse than random guessing (AUC < 0.5, Fig 1A), suggesting that the current incomplete knowledge of virus diversity could lead to entirely incorrect priorities under such approaches. In contrast, models that exploited features of viral genomes that were at least partly independent of virus taxonomy both generalised predictions across divergent viruses and provided capacity to discriminate risk among closely related virus species.

In requiring only a genome sequence, our approach has quantitative and qualitative advantages over alternative models for zoonotic risk assessment. The most comprehensive alternative model requires virus species-level information on publication count (a proxy for study effort), the diversity of hosts infected, whether or not the virus is vector borne, and the ability to replicate in the cytoplasm [5]. We estimate similar predictive performance for this model (AUC_m_ = 0.770) using a subset of only mammalian viruses (see methods). However, neither study effort nor knowledge of host range are available for novel viruses, and restricting this model to factors which might reasonably be inferred from virus taxonomy (vector borne status and ability to replicate in the cytoplasm) performs considerably worse (AUC_m_ = 0.647) than our approach. We were unable to compare the performance of our model to a more recently developed prioritisation system based on ecological variables weighted by expert opinion, as metrics of performance were not provided and could not be calculated for a comparable set of viruses with known zoonotic status [6]. Although we emphasize that the viruses included and study objectives differed, that genome-based ranking seems to perform comparably to or better than currently-available alternatives that require far more, and often unavailable, data highlights the surprisingly informative signals of human infection-ability contained within viral genomes. Crucially, from the perspective of virus risk assessment, models based purely on genome sequences can be applied much earlier to identify many potential zoonoses immediately after virus discovery and genome sequencing, when data on most other risk factors are still unknown. Ultimately, genome-based rankings could be combined with data on additional known risk factors as they become available [5,6].

By highlighting viruses with the greatest potential to become zoonotic, genome-based ranking allows further ecological and virological characterisation to be targeted more effectively. Indeed, studying viruses in the order suggested by genome-based ranking would find many zoonotic viruses much earlier than current taxonomic or phylogenetically-informed approaches (Fig 1D & S6 Fig). Nevertheless, we acknowledge that even after applying our models, considerable numbers of viruses may need to undergo confirmatory testing (e.g. infectivity experiments on human-derived cell lines [23]) before significant further research investments, and this need will only increase with ongoing virus discovery. Although these numbers are more manageable considering that experimental validation will be dispersed across virus taxonomic groups which will be studied by different experts, efforts to increase the success rates of virus isolation (a prerequisite for current *in vitro* validation methods) and to create systems for high-throughput virus host range testing are clearly needed to improve the efficiency of this process [23,24]. Such efforts could further generate valuable feedback data; iteratively improving model performance and consequently reducing the relative proportion of new viruses requiring additional laboratory testing.

Several lines of evidence – including SHAP clustering of viruses with different genome organizations, accurate prediction of human-infecting viruses from families withheld from training, and the prediction of the zoonotic risk of SARS-CoV-2 when withholding data from other zoonotic sarbecoviruses – suggested that our models make predictions using genomic features that predict human infection across divergent virus taxa. From a practical standpoint, this is a major advantage since it means that our model borrows information across families and might therefore anticipate the zoonotic potential of viruses which, due to their rarity or lack of historical precedent, would not otherwise be considered high risk (sometimes referred to as Disease X) [25]. From a broader evolutionary standpoint, the putative existence of convergently evolved features in viral genomes which seem to predispose human infection is a discovery which deserves further mechanistic study. Encouragingly, a substantial literature in vaccine development facilitates genome-wide synonymous recoding of viral genome composition and has established that these changes can dramatically affect viral fitness [26,27]. Our results provide a path by which analogous approaches could test how the features we identified affect viral host range in general and human infection ability in particular. Doing so may reveal novel mechanisms of viral adaptation to humans, which might represent both biologically-verified risk factors for the improvement of future models of zoonotic potential and potential therapeutic targets.

We used single exemplar genomes from each virus species to maximise the likelihood of discovering generalisable signatures of human infection while avoiding performance measures which would be over-optimistic for novel viruses. A potential drawback of this approach was that we omitted substantial viral diversity which is not yet formally recognized by the ICTV [3]. However, we contend that including currently unrecognized viruses is unlikely to improve the predictions of our models because: (a) most will be non-human infecting (an already over-represented class) and hence provide little additional information, (b) those which do infect humans will not generally be known to do so due to a lack of historic testing, adding misleading signals, and (c) the predictive features identified often span across families, reducing the impacts of taxonomic gaps. The use of single genomes does however mean that the ranks produced here pertain only to the specific genomes tested (in most cases, the NCBI reference sequence, S1 Table), and may not apply equally across all strains within a species (Fig 5B). We also stress that our model predicts baseline zoonotic *potential* (i.e., ability to infect humans), which ultimately will be modulated by ecological opportunities for emergence [28,29]. Further, the societal impact of emergence will depend on capacity for human to human spread and on the severity of human disease, which likely require additional non-genomic data to anticipate [28,30].

In summary, we have constructed a genomic model that can retrospectively or prospectively predict the probability that viruses will be able to infect humans. The success of our models required aspects of genome composition calculated both directly from viral genomes and in units of similarity to human transcripts, and some viruses were predicted to be zoonotic due to common genomic traits despite ancient evolutionary divergence. This highlights the potential existence of currently unknown phenotypic consequences of viral genome composition that appear to influence viral host range across divergent viral families. Independently of the mechanisms involved, the performance of our models shows how increasingly ubiquitous and low-cost genome sequence data can inform decisions on virus research and surveillance priorities at the earliest stage of virus discovery with virtually no extra financial or time investment.

## Materials and methods

### Data

Although our primary interest was in zoonotic transmission, we trained models to predict the ability to infect humans in general, reasoning that patterns found in viruses predominately maintained by human-to-human transmission may contain genomic signals which also apply to zoonotic viruses. Data on the ability to infect humans were obtained by merging the data of [5], [9], and [17], which contain species-level records of reported human infections, resulting in a final dataset of 861 virus species from 36 families. In all cases, only viruses detected in humans by either PCR or sequencing were considered to have proven ability to infect humans. All viruses for which no such reports were found were considered to not infect humans (as long as they were assessed for potential human infection by at least one of the studies above), although we emphasise that many of these viruses are poorly characterised and could therefore be unrecognized or unreported zoonoses. We therefore expect our models to further improve as these and new viruses become better characterised. For figures, human-infecting viruses were further separated into primarily human-transmitted viruses and zoonotic viruses, based on virus reservoirs recorded in [9]. Human-infecting viruses for which the reservoir remains unknown were assumed to be zoonotic, while viruses with both a human and non-human reservoir cycle (e.g., *Dengue virus*) were recorded as primarily human-transmitted, reflecting the primary source of human infection. A representative genome was selected for each virus species, giving preference to sequences from the RefSeq database wherever possible. RefSeq sequences that had annotation issues, represented extensively passaged isolates, or were otherwise not judged to be representative of the naturally circulating virus were replaced with alternative genomes.

### Features

We compared the predictive power of all classifiers to that which could be obtained through knowledge of a virus’ taxonomic position alone. This captures the intuitive expectation that viruses can be risk-assessed based on knowledge of the human-infection abilities of their closest known relatives. To formalise this idea, we first created a simple heuristic which ranks viruses based on the proportion of other viruses in the same family which are known to infect humans (‘*taxonomy-based heuristic*’ in Fig 1). Viral family was chosen as the level of comparison because not all viruses are classified in a scheme which includes subfamilies, while lower taxonomic levels suffer from limited sample size. To further characterise the predictive power of virus taxonomy, we also included potential predictor variables (here termed *features*) describing virus taxonomy when training classifiers. These included the proportion of human infecting viruses in each family (calculated from species in the training data only) along with categorical features describing the phylum, subphylum, class, subclass, order, suborder, and family to which each species was assigned (‘*taxonomic feature set’*, 8 features). This information was taken from version 2018b of the ICTV master species list (https://talk.ictvonline.org/files/master-species-lists/). To capture taxonomic effects at finer resolution, we summarised the human infection ability of the closest relatives of each virus in the training data (following [10], here termed the ‘*phylogenetic neighbourhood feature set*’, 2 features). To calculate these features, the genome (or genome segments, where applicable) of each virus species was nucleotide BLASTed against a database containing genomes or genome segments for all species in the training data. All BLAST matches with e-value ≤ 0.001 were retained and used to calculate the proportion of human-infecting viruses in the phylogenetic neighbourhood of each virus (excluding the current species). We also calculated a ‘distance-corrected’ version of this proportion by re-weighting matches according to the proportion of nucleotides *p* matching the genome of the focal virus:

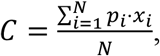

where *N* is the number of retained blast matches for the focal virus species, and *x_i_* = 1 if matching species *i* is able to infect humans and equals 0 otherwise. Both the raw and distance-corrected proportion were used in unison to define the phylogenetic neighbourhood, allowing classifiers to pick the most informative representation or to combine both pieces of information if needed. In cases where a virus received no matches with e-value ≤ 0.001, both proportions were set to NA to reflect the fact that no information about the phylogenetic neighbourhood was available. This occurred for an average of 2% of viruses in each random training and test set (see below).

Various features summarising the compositional biases in each virus genome were calculated as described in [10]. These included codon usage biases, amino acid biases, dinucleotide biases across the entire genome, dinucleotide biases across coding regions only, dinucleotide biases spanning the bridges between codons (i.e. across base 3 of the preceding codon and base 1 of the current codon), and dinucleotide biases at non-bridge positions (‘*viral genomic features*’, 146 features). Similarity features to human RNA transcripts were obtained by first calculating the above compositional biases for human genes. For each gene, the sequence of the canonical transcript was obtained from version 96 of Ensembl [31]. Genes were divided into three mutually exclusive sets, encompassing interferon stimulated genes (ISGs, taken from [13]; N = 2054), non-ISG housekeeping genes ([32]; N=3172), and remaining genes (N=9565). The distribution of observed values for each genome feature was summarised across all genes in a set by calculating an empirical probability density function using version 2.3.1 of the EnvStats library in R version 3.5.1 [33]. The final similarity score for each genome feature of each virus was then calculated by evaluating this density function at the value observed in the virus genome, giving the probability of observing this value among the transcripts of the set of human genes in question (S12 Fig). This yielded three feature sets termed ‘*similarity to ISGs*’, ‘*similarity to housekeeping genes*’ and ‘*similarity to remaining genes*’, each containing 146 features.

### Training

Gradient boosted classification trees were trained using the xgboost and caret libraries (versions 0.90 and 6.0-85, respectively) in R [34,35]. Note that while we separate primarily human-transmitted viruses and zoonotic viruses in some figures, these viruses were considered a single class during training. Thus, models were trained to distinguish viruses known to infect humans from those with no reports of human infection.

A range of models were trained using different combinations of the individual feature sets described above. To reduce run-times and potential overfitting in subsequent steps, features were subjected to a pre-screening step to remove those with little or no predictive value. During this pre-screening step, models were trained on a random selection of 70% of the data using all features within the respective feature group (e.g., all ISG similarity features). Training sets were selected using stratified sampling (i.e., selecting positive and negative examples separately) to retain the observed frequencies of positive (known human-infecting) and negative (not known to infect humans) virus species. We were unable to additionally stratify training set selection by virus family due to the small numbers of species in many families. All hyperparameters were kept at their default values, except for the number of training rounds, which was fixed at 150. The importance of each feature was summarised across 100 iterations in which the same features were used to train a model on different samples of the full dataset, and the *N* most predictive features were retained. A range of possible values for *N* was evaluated by combining all feature sets and using the selected features to optimize and train a final set of model iterations as described below. The final value of *N* = 125 was chosen as the point at which additional features provided no further improvement in performance, measured as the area under the receiver-operating characteristic curve (AUC; S13 Fig). Here, AUC measures the probability that a randomly chosen human-infecting virus would be ranked higher than a randomly chosen virus which has not been reported to infect humans. When a given feature set or combination of feature sets comprised < 125 features, all features were retained.

Final models were trained using reduced feature sets. To assess the variability in accuracy across different training sets, training was repeated 100 times [10]. In each iteration, training was performed on a random, class-stratified selection of 70% of the available data (here, the *training set*). Output probabilities were calibrated using half the remaining data (*calibration set*, again selected randomly and stratified by human infection status), leaving 15% of the full dataset for evaluation of model predictions (*test set*). In each iteration, hyperparameters were selected using 5-fold cross-validation on the training set, searching across a random grid of 500 hyperparameter combinations. This cross-validation was adaptive, evaluating each parameter combination on a minimum of 3 folds before continuing cross-validation only with the most promising candidates [35]. The parameter combination maximising AUC across folds was selected and used to train a final model on the entire training set. This model was then used to produce quantitative scores for each species in the calibration and test sets.

Next, outputs were calibrated to allow interpretation as probabilities using the beta calibration method of [36]. A calibration model was fit to scores obtained for the calibration set using version 0.1.0 of the betacal R package. The fitted calibration model was used to produce final output probabilities for virus species in the test set. Finally, to summarise predicted probabilities from the same model trained on different training sets, we averaged the calibrated probabilities across the best-performing iterations (a process with similarities to bagging [10]; 1000 iterations performed). Bagging of virus species from the training data relied on the best 10% of iterations in which each virus occurred in the test set. As such, the focal virus had no influence on the training or calibration of the iterations used in bagging. Further, the performance of each model iteration was re-calculated while excluding the focal species from the test set, to prevent accurate prediction of the focal virus from influencing the choice of iterations used for bagging. When predicting the probability of human infectivity for viruses that were completely separated from training (i.e., those in our first case study), we used the best 10% of iterations overall.

Calculating phylogenetic neighbourhood features represented a bottleneck in the iterative training strategy described above, because each iteration had a different reference BLAST database, corresponding to the specific training set selected. This would have required repeating BLAST searches at every iteration. To overcome this, a single all-against-all blast search was performed, with search results and e-values subsequently corrected in each iteration to emulate the result that would have been obtained when blasting only against the current training dataset. Specifically, e-values were re-calculated as described in [37]:

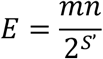

where *m* is the length of the query sequence (in nucleotides), *n* is the total number of nucleotides in the training set (i.e., the size of the database searched), and *S*’ is bitscore for this particular alignment in the original blast search.

### Feature importance and clustering

To assess the variability in feature importance while accounting for all viruses, feature importance was assessed across all 1000 iterations produced for bagging above. In each iteration, the influence of features was assessed using SHAP values, an approximation of Shapley values which here describe the change in the predicted log odds of infecting humans attributable to each genome composition feature used in the final model [18]. In each iteration, this produced a SHAP value for each virus–feature combination. The overall importance of each feature was calculated as the mean of absolute SHAP values across all viruses in the training set of a given iteration [38].

Because features tended to be highly correlated, we also report importance values for clusters of correlated features, with the importance of each cluster for individual viruses calculated as the sum of absolute SHAP values across all features in a cluster:

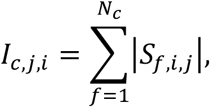

where *I_c,j,i_* is the importance of feature cluster *c* in determining the output score of virus *j* in iteration *i*, *N_c_* is the number of features in this cluster, and *S_f,i,j_* is the SHAP value for feature *f*. The overall importance of each feature cluster in a given iteration was then calculated as the mean of these importance values across all viruses in the training data of that iteration:

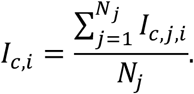

Feature clusters were obtained by affinity propagation clustering, which seeks to identify discrete clusters of features centred around a representative feature (the *exemplar* feature) [39]. Features were clustered using pairwise Spearman correlations as the similarity measure, using version 1.4.8 of the apcluster library in R [40].

To further explore patterns in feature importance across virus species, we followed a strategy similar to [38], clustering viruses based on the average SHAP values assigned to individual features for each virus across all iterations:

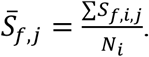

These values were used to calculate the pairwise Euclidean distances between all virus species using version 2.1.0 of the cluster library in R [41]. Viruses were then clustered using agglomerative hierarchical clustering, calculating distances between clusters as the mean distance between all points in the respective clusters (i.e., UPGMA clustering). To explore patterns common to viruses from each class, clustering was performed separately for known human infecting and other viruses.

To compare this explanation-based clustering with virus taxonomy, we also constructed a dendrogram based on taxonomic assignments as recorded in version 2018b of the ICTV master species list, using all taxonomic levels from phylum to subgenus. Since some levels of the ICTV taxonomy are not used consistently across all viruses, missing taxonomic levels were interpolated to ensure accurate representation of the underlying taxonomy. For example, for viruses which are not classified in a scheme which includes subfamilies, the next level downstream – genus – was repeated, thereby treating each genus as belonging to a distinct subfamily. Categorical taxonomic assignments were used to calculate pairwise Gower distances between virus species [42], before performing agglomerative hierarchical clustering as described above. We also assessed the ability of underlying genome feature values to reconstruct virus taxonomy by performing hierarchical clustering on a Euclidean distance matrix calculated directly from all genome composition features (i.e., the unreferenced genome, ISG similarity, housekeeping gene similarity and remaining gene similarity feature sets). The similarity between dendrograms was assessed using the gamma correlation index of [19], as implemented in dendextend version 1.12.0 in R [43]. A null distribution for this statistic was calculated by randomly shuffling the labels (i.e., virus species names) of both dendrograms 1000 times. To assess the taxonomic depth at which dendrograms were concordant, the Fowlkes-Mallows index was calculated at each possible cut-point in the dendrograms being compared [44], again using the dendextend library. As before, a null distribution was generated by randomly shuffling the labels of both dendrograms 1000 times.

### Ranking holdout viruses

To illustrate the use of our models in practice, the best performing model (i.e. the bagged model trained using the best 125 features selected from among all genome composition-based feature sets, here termed the ‘*combined genome feature-based model*’) was used to generate predictions for a set of held out viruses. We included all virus species recognized in the latest version of the ICTV taxonomy (release #35, 2019; https://talk.ictvonline.org/taxonomy) which were from families known to contain species that infect animals but which did not occur in our training data because they were absent from the previously described databases of human infection ability used to form the training data [5,9,17]. These included all 36 families represented in the training data, plus *Anelloviridae* and *Genomoviridae*. Names of viruses in the training data were updated to the latest taxonomy and checked for matches in the corresponding ICTV master species list before extracting non-matching species. For each species, the genome sequence referenced in the ICTV virus metadata resource corresponding to this version of the taxonomy (https://talk.ictvonline.org/taxonomy/vmr) was retrieved and used to calculate genome composition features as described above. The host from which each virus genome was generated was obtained from either the corresponding GenBank entry, the publication first describing the sequence, or the ArboCat database. This host information was used to further subset viruses to include only those sampled from birds, mammals, Diptera (which includes common vectors such as mosquitos and sandflies) and Ixodida (ticks), or for which the sampled host could not be identified. Scoring was performed using each of the top 10% out of 1000 iterations, as described above, and averaged to obtain the final output probability. Confidence intervals for these mean probabilities were calculated as the 2.5% and 97.5% quantiles of probabilities output by the top 10% of classifiers. Tests for the effects of virus family and sampled host on the predicted probability of infecting humans were performed by fitting a beta regression model using version 1.8-27 of the mgcv library in R. This model fitted the mean predicted probability for each virus as a function of whether a human host was sampled, whether the sample derived from arthropods (both binary fixed effects), random effects for the taxonomic order and class of sampled hosts, and a random effect for virus family. Partial effects plots were generated using code from [9].

### Evaluating existing model performance

To compare the performance of our models to previously published ecological models, we downloaded the fitted models of [5]. The best viral traits model fitted while excluding serological detections (termed “stringent data” in [5], N = 408) was then subjected to a testing regime similar to that used for our models. Across 100 iterations, models were re-fit using a randomly selected subset consisting of 85% of the virus species used in [5]. Each fitted model was then used to predict probabilities of being zoonotic for the remaining 15% of species. This matched the evaluation strategy used for our own models (as reported in Fig 1A), with 85% of the data used during training and calibration, leaving 15% of the data to test model accuracy. Predictions from each iteration were used to calculate AUC using version 1.2.0 of the ModelMetrics package in R.

## Acknowledgments

We thank Laura Bergner, Richard Orton, Andrew Shaw, Sam Wilson and David Robertson for helpful discussions and suggestions. D.G.S. and N.M. were supported by a Wellcome Senior Research Fellowship (217221/Z/19/Z). Additional funding was provided by the Medical Research Council through program grants MC_UU_12014/8 and MC_UU_12014/12. The funders had no role in study design, data collection and analysis, decision to publish, or preparation of the manuscript.

## Data and materials availability

Data and code related to this paper are available at: https://doi.org/10.5281/zenodo.4271479

