## Supplementary material for "Identifying and prioritizing potential human-infecting viruses from their genome sequences"

**This PDF file includes:**

Supplementary text (S1 Text)
Supplementary figures (S1 Fig – S13 Fig)

**Other Supplementary Materials:**

S1 Table. (separate file)

**S1 Text: Viral genome compositional predictors of human infection.**

Our results showed that combining information on a variety of genome compositional biases can differentiate viruses known to infect humans from those which have not been reported to do so. Moreover, the aspects of genome composition which predicted human infection in some cases did so across divergent viruses (e.g., from different taxonomic classes), signalling potential for convergent evolutionary processes which predispose the capability for human infection through mechanisms of virus-host adaptation which are not currently understood. We therefore carried out additional analyses exploring the influence of our input genome composition measures on model predictions in order to aid future research using experimental approaches to manipulate viral genome composition.

First, we compared the frequency at which features calculated directly from the viral genome or in reference to three sets of human transcripts were included in the final model. We hypothesized that human similarity features would be more prominent if viral mimicry of host transcripts increased viral fitness in humans (either due to common selective pressures from nucleotide sensing defences or enhanced translation) or if dissimilarity to human transcripts was somehow adaptive. In contrast, unreferenced viral genomic features might be more prominent if specific aspects of genome composition were adaptive for viruses, but neutral for human genes. Among the 125 features retained in the final model, most described similarity of viral genomic features to the transcripts of human genes (ISGs = 27.3%; housekeeping genes = 22%; remaining genes = 12.7%). The remaining 38% of features described viral genomic composition directly. Although this excess frequency of human similarity-based features among retained features is partly explained by the larger number of such features available, compositional similarity to housekeeping genes and ISGs also tended to have a larger magnitude of influence on predictions than unreferenced viral genomic features (Fig 2B). Moreover, when different representations of the same genome composition measure were retained in the final model (N = 20 compositional measures, 50 features), the human similarity version tended to be more important than the unreferenced measure of viral genomic composition (Fig 2C). These results imply that when retained, features that described similarity to human genes had more powerful effects on model predictions of human infection.

Since many genome compositional features were correlated within viral genomes, we further summarised the total predictive power of 31 discrete clusters of features identified by affinity propagation clustering of pairwise Spearman correlations between observed feature values (Fig 2D & S9 – S10 Fig). As expected, many clusters (41.9%) contained a mixture of different feature types (amino acid biases, codon biases, and dinucleotide biases), with 4 clusters containing combinations of all three. As a result, no single genome composition measure could be isolated as the driving force behind the ability of models to predict human infection ability. Most clusters also contained both unreferenced viral genomic and similarity-based features (Fig 2D), but ISG similarity features were identified as the exemplars (i.e., the most central or representative datapoint) of the largest number of clusters consisting of ≥ 3 features (36%, observed/expected ratio [OER] = 1.40). Housekeeping and remaining gene similarity features were identified as the exemplars of 24% and 16% of such clusters, respectively (OER = 1.08 and 1.64), leaving unreferenced genome features under-represented as exemplars (24%, OER = 0.57). Measures of similarity to human genes were therefore the best representation of the majority of feature clusters predicting human infection ability (Fig 2D & S9 Fig).

Acknowledging the challenge in interpreting correlated genomic features in isolation, we next explored the shape of relationships between the values of each feature and the predicted log odds of human infection (S10 Fig). For human similarity features, we observed patterns consistent with both compositional mimicry and, more rarely, compositional distancing. For both positive and negative effects, patterns could be linear or non-linear (e.g., extreme values increasing the odds of human infection, but all others being neutral or negative; see for example similarity in TpC-dinucleotide bias at codon bridges [brTpC similarity]). Similarly, for unreferenced viral genomic features, we observe both positive (e.g., GpT-dinucleotide bias and serine amino acid bias) and negative (e.g., leucine amino acid bias and GpA dinucleotide bias at codon bridges) relationships with human infection, implying that both avoidance and favouring of certain genomic features can increase the odds of human infection. This finding in part arises because many of the genome compositional features we included are (or form part of) redundancies in the genetic code, such that bias against one feature necessarily implies bias for another, along with uncertainty surrounding which parts of the human genome might be mimicked. For example, high levels of brTpC similarity to remaining genes was associated with increased likelihood of infecting humans, but high levels of similarity in the same feature to the transcripts of housekeeping genes had the opposite effect (S10 Fig). However, these two measures of similarity were correlated (Spearman correlation = 0.53) and were grouped in the same cluster of features, with brTpC similarity to the transcripts of remaining genes as the exemplar (cluster 2, S9 Fig). The transcripts of remaining genes tended to have a slightly lower TpC bias at codon bridges (median = 0.95) than those of housekeeping genes (median = 1.03). Together, these observations suggest that if mimicry of bridge TpC bias is the factor under selection (as opposed to any of the other correlated features in cluster 2) – it appears that matching remaining genes is more important, while the measured similarity to housekeeping genes may simply act as a marker for too-high TpC content. Similar patterns were observed across the majority of clusters, with at least one of the correlated features in each of these clusters supporting a role for mimicry of human gene transcripts. However, three clusters stood out as pointing to a negative relationship between similarity and human infection ability without any correlated features supporting a role for mimicry (clusters 14, 20, and 26, S9 Fig). These apparent compositional distancing effects tended to have weak effects compared to compositional mimicry features and the mechanisms driving their association with human infection are unknown. It is possible that distancing reflects complex trade-offs between different features or mimicry of a set of human gene transcripts that is poorly captured by the three gene sets summarised here. Alternatively, apparent compositional distancing may arise from selection to avoid specific features at all costs (e.g. those targeted by antiviral defences), where a low occurrence of the feature may be more important than matching the particular frequency found in human genes subject to additional constraints. Interestingly, several features strongly linked to human infection here were also identified among the most powerful features for discriminating reservoir hosts in a smaller dataset of RNA viruses, including leucine and serine amino acid bias and GpT-dinucleotide bias, supporting their potential role in influencing viral host range (see figure S10 in [1]).

Taken together, these results suggest that matching (or, potentially, diverging from) human genes either serendipitously or by selection in natural host species, along with other mechanisms by which viruses adapt their genome composition in ways that are independent of host genome composition, predictably predisposes some viruses to be capable of infecting humans. That viruses exploit a diversity of approaches is intuitive since our dataset contained a diverse array of RNA and DNA viruses which are likely both to face different selective pressures and to employ different solutions in the case of common selective pressures. Even within a single virus species, a combination of effects which involve, for example, avoidance of innate immunity by reducing certain dinucleotide motifs and mimicry of other motifs which enhance translation could together enhance the likelihood of human infection. Although the exact mechanisms by which genome compositional biases affect viral fitness are only beginning to be understood, the existence of these fitness effects, including effects on host range, are increasingly evident for a variety of viruses [2–5]. Experimental approaches involving viral genome re-coding are well developed from the virus attenuation literature [reviewed in 6] and would be useful to isolate the effects of individual aspects of genome composition suggested by our predictive model.

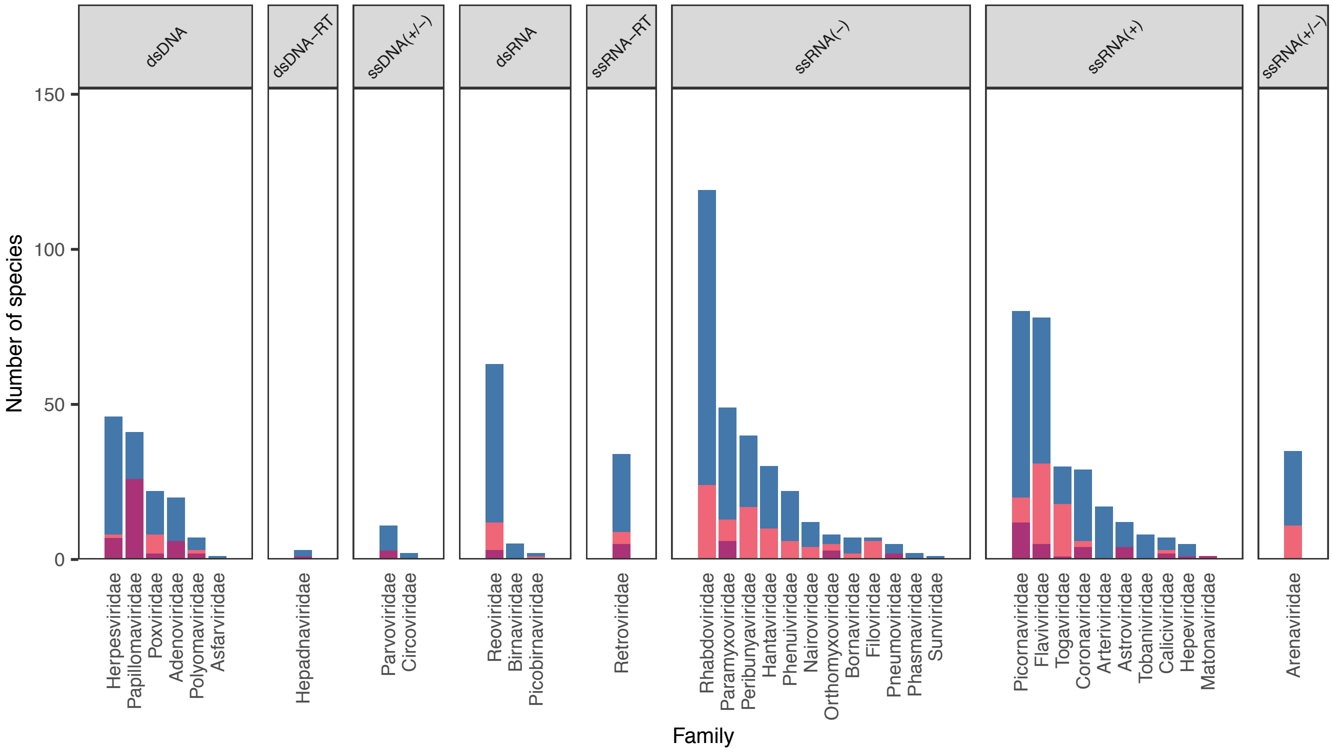


**S1 Fig. Data used in this study for model development and evaluation.**
Viruses (N = 861 species) predominately transmitted among humans (purple) or from animals to humans (zoonoses, pink) were combined to form the positive class of human-infecting viruses. All other virus species, for which no human infections have been detected (blue), were used as the negative class when training models.

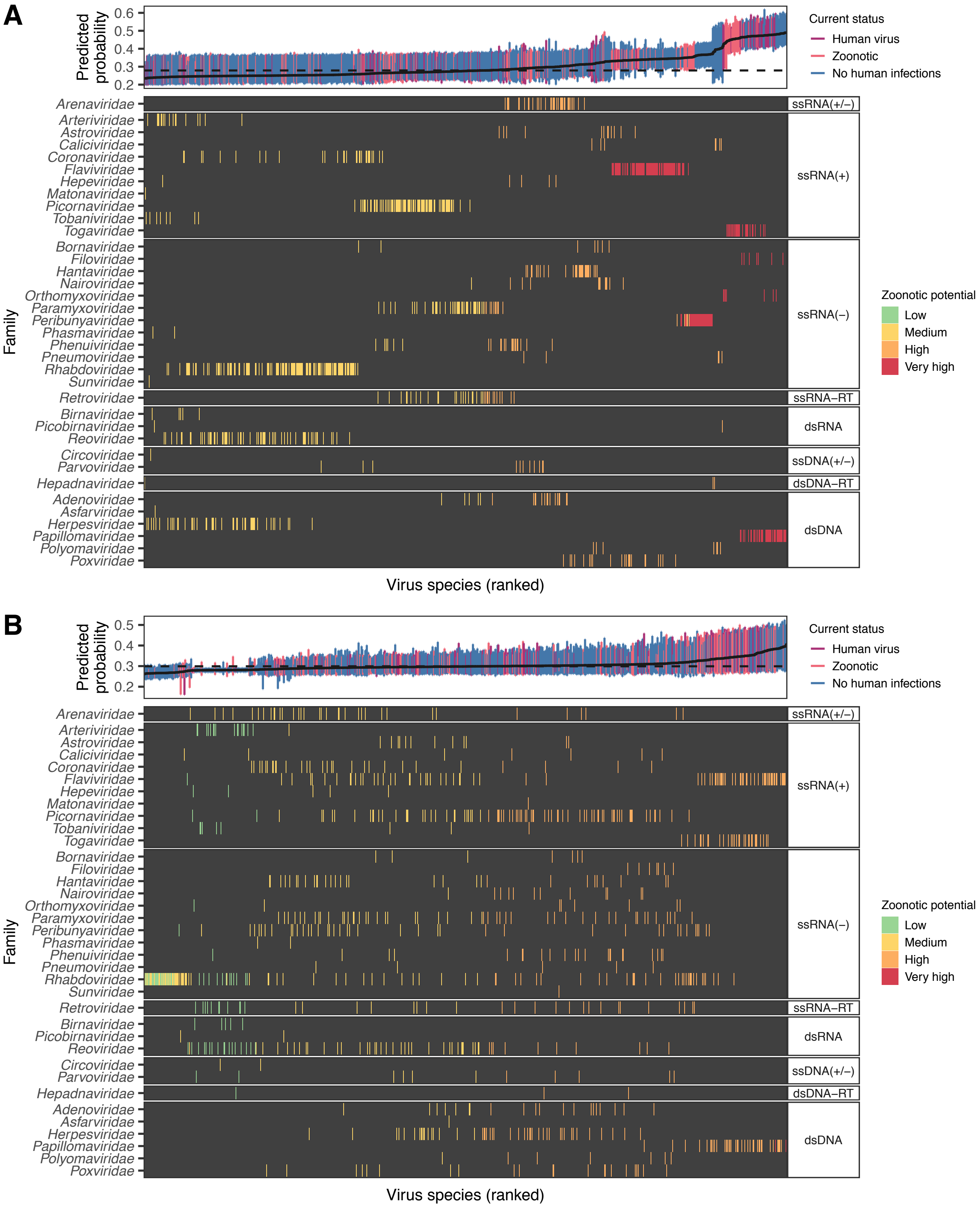


**S2 Fig. Virus ranks produced by relatedness-based models have limited ability to discriminate the zoonotic potential of closely-related viruses.**

Viruses from the training data are shown ranked by their predicted probability of infecting humans produced by bagged versions of (A) the taxonomic feature set-based model and (B) the phylogenetic neighbourhood (PN)-based model. In the top panel of each plot, a grey line shows mean bagged probabilities, while coloured error bars highlight the region containing 95% of predictions from the models used in bagging. A dashed line shows the cutoff which balances sensitivity and specificity. The lower panel in each plot highlights the location of viruses from different families (coloured bars) and contains a dark grey background for clarity. Clustering of zoonotic risk by virus family in the taxonomic features model (A, lower panel) shows that this model is unable to discriminate high and low risk viral taxa within viral families. This was expected, since this model contained only features resolved at the family level or above. A sequence similarity-based approach was expected to perform better, but while the PN model (B) does show improved resolution, it still fails to separate risk within certain viral families. For example, papillomaviruses and most flaviruses are ranked as high risk, despite the presence of non-zoonotic species within each family. In contrast, the genome composition model was considerably more accurate (Fig 1 and S3 Fig) and was able to assign viruses from the same family into a broader range of risk categories (see S4 Fig), consistent with our current biological understanding that zoonotic ability varies within viral families.

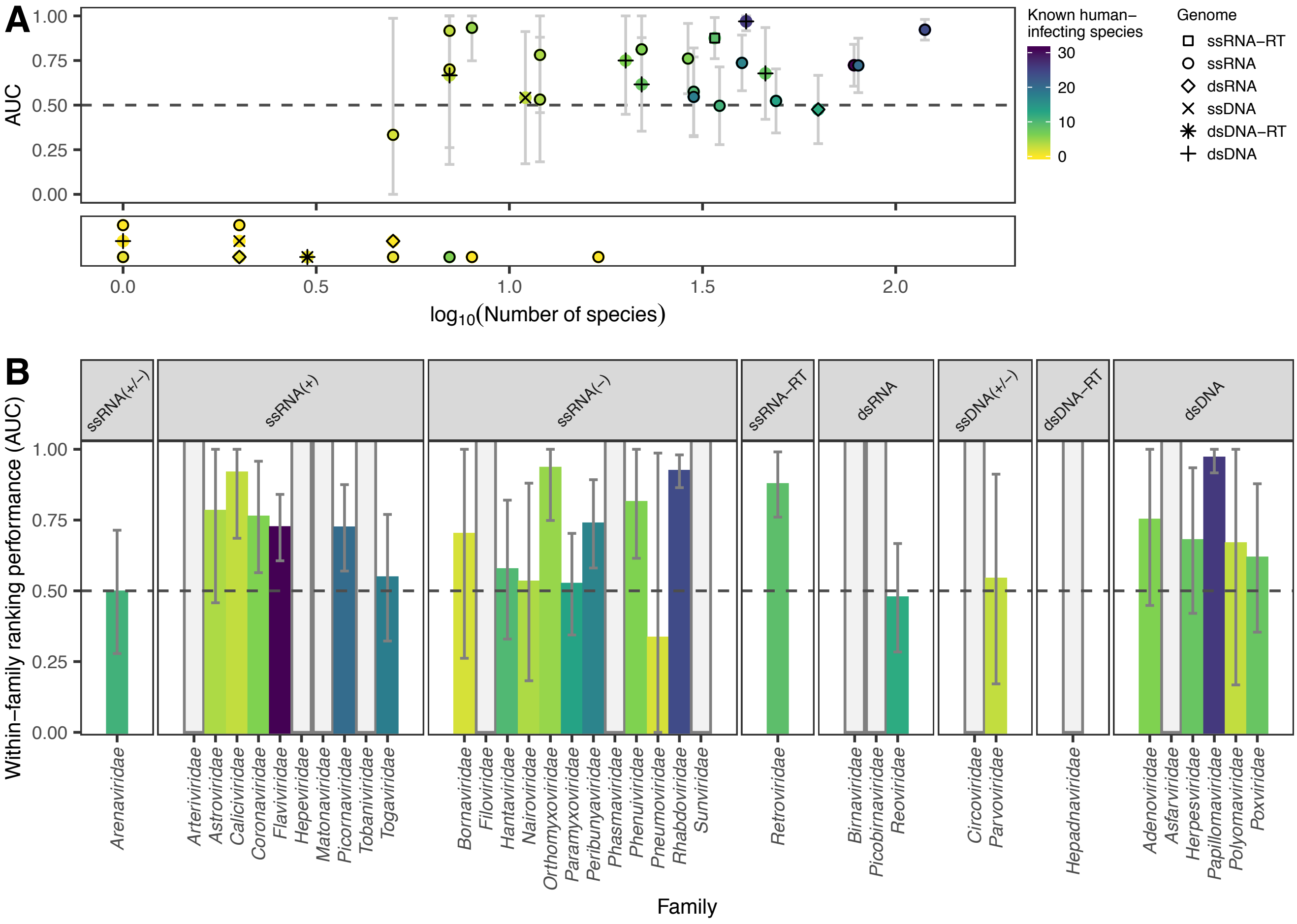


**S3 Fig. Family-specific measures of accuracy for the combined genome feature-based model.**

AUC here measures the probability of accurately ranking known human-infecting viruses above other viruses from the same family, when ranking viruses using the output from the bagged model based on all genome feature sets. AUC values could not be calculated for families containing < 2 human infecting viruses or < 2 viruses not known to infect humans. These families are illustrated in the lower plot in (A), where the y-axis is unitless and overlapping points are stacked, and as grey bars in (B). Confidence intervals were calculated using the method of [7,8], and highlight the difficulty of assessing within-family AUC given the relatively small numbers of viruses currently known in most families. We detected no obvious taxonomic pattern in the variation of within-family AUC values.

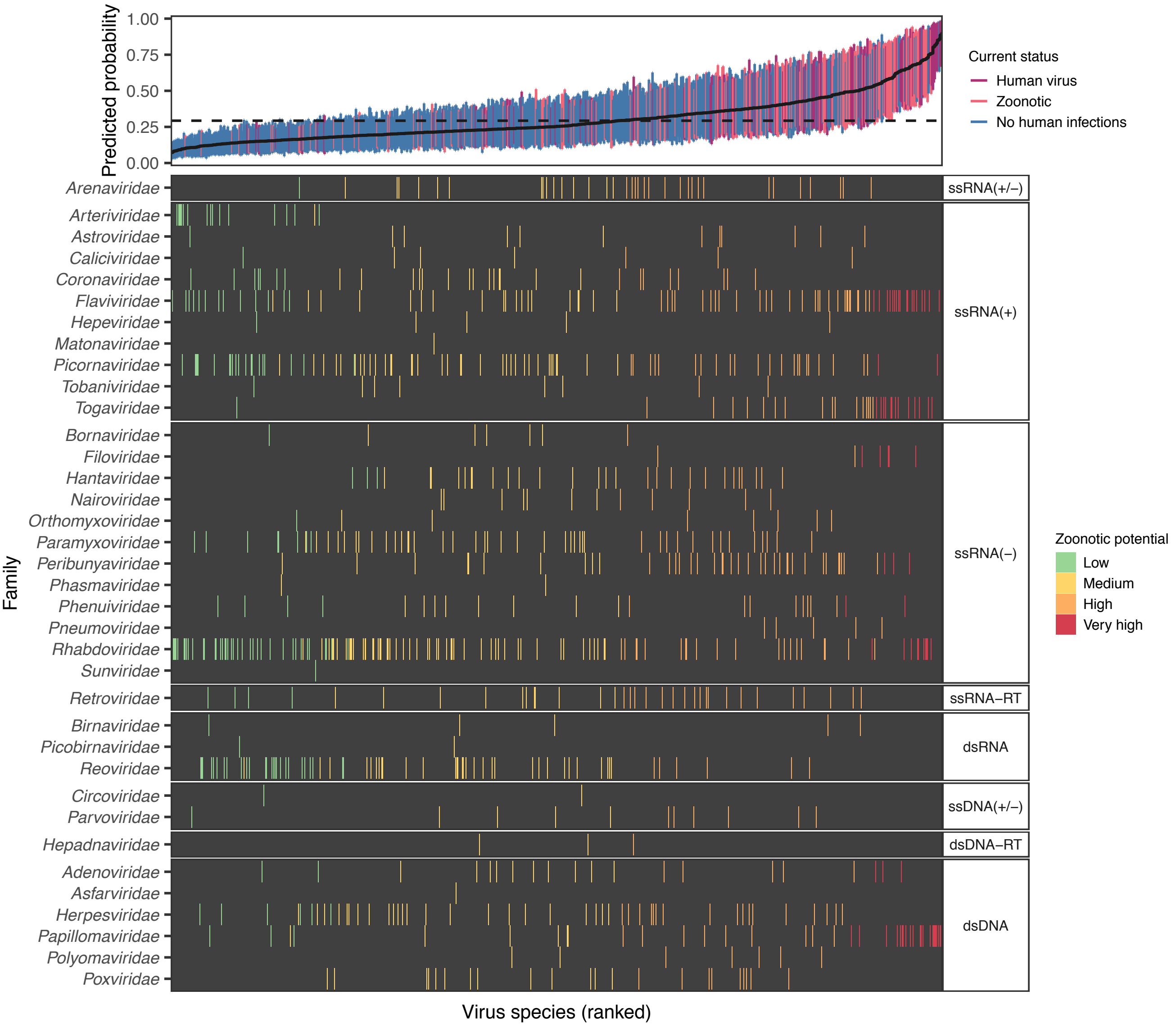


**S4 Fig. Heterogeneous zoonotic risk predictions for species within viral families.** Viruses in the training data are shown ranked by their mean predicted probability of infecting humans, produced by bagging across iterations of the combined genome feature-based model. In the top panel, a grey line shows mean bagged probabilities, while coloured error bars highlight the region containing 95% of predictions from the models used in bagging. A dashed line shows the cutoff which balances sensitivity and specificity. The lower panel highlights the location of viruses from each family using coloured bars, and contains a dark grey background for clarity. For a detailed list of viruses and their ranks and priorities, see S1 Table. Model predictions within viral families span risk categories, illustrating the power to discriminate risk at higher taxonomic resolution than models based on conserved features of virus biology (e.g. ability to replicate in cytoplasm or be transmitted by arthropod vectors), or alternative models based on taxonomy or phylogenetic neighbourhood (S2 Fig).

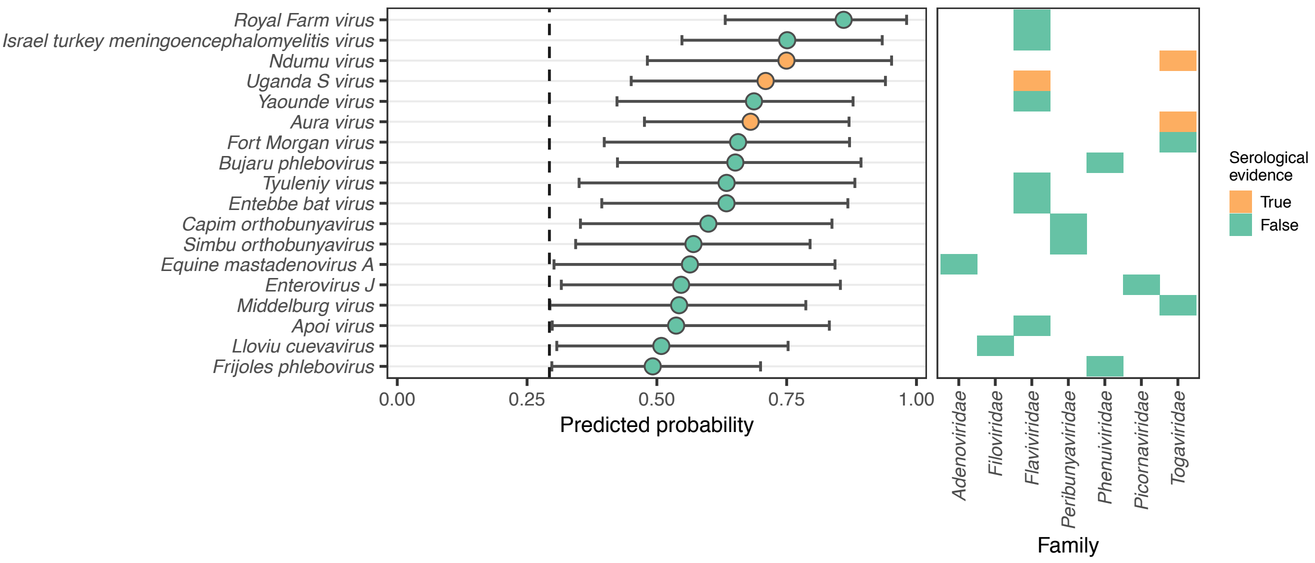


**S5 Fig. Putative unrecognized zoonoses identified within the training data.**

Points and confidence intervals in the left panel show the bagged mean and 95% confidence range on the predicted probability of human infection when using the combined genome feature-based model. Each virus species shown was included in the training data as not currently known to infect humans, but was nevertheless classified in the 'very high zoonotic potential' category when included in test sets, which means ≥ 95% of the bagged models predict these viruses as human-infecting. The right panel shows evidence of serological infection for three of these viruses (orange), from [9,10]. Absence of serological evidence may reflect lack of diagnostic testing, lack of quantifiable antibody responses, lack of human exposures or inability to infect humans.

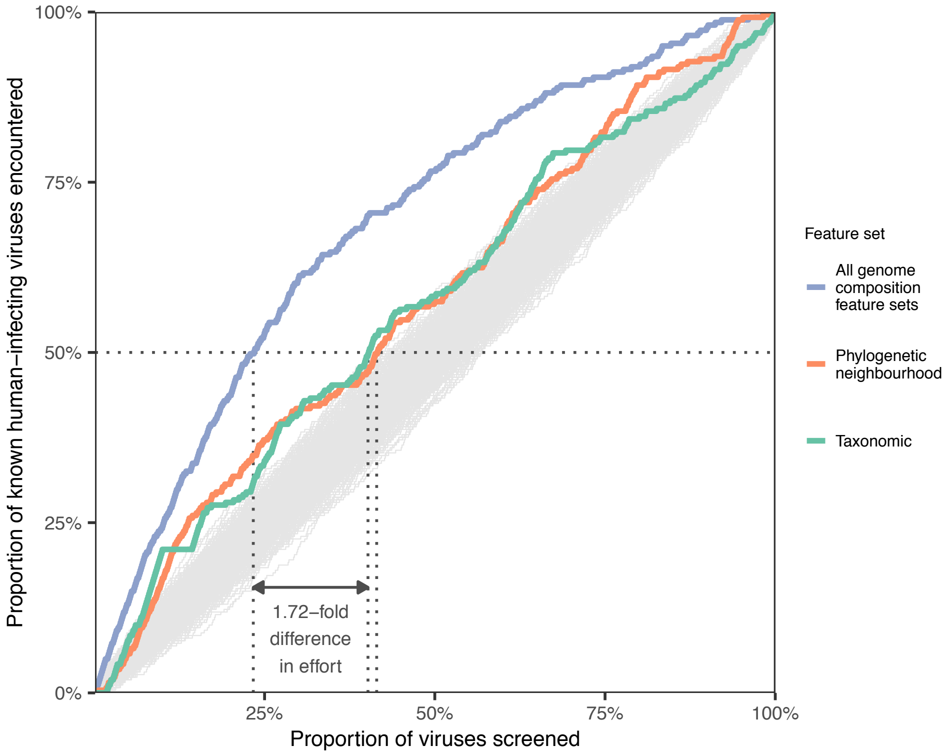


**S6 Fig. Cumulative evaluation of human-infecting species when viruses are prioritised for downstream research or surveillance in the order suggested by models trained on different feature sets.**

For each feature set (or combination of feature sets), viruses were ranked based on bagged predictions across the top 100 out of 1000 training iterations. Dotted lines highlight the proportion of all viruses in the training and evaluation data which need to be screened to detect 50% of known human-infecting viruses. Grey lines show the range of accumulation curves expected from random screening in a dataset of this size, simulated by randomly shuffling viruses 1000 times.

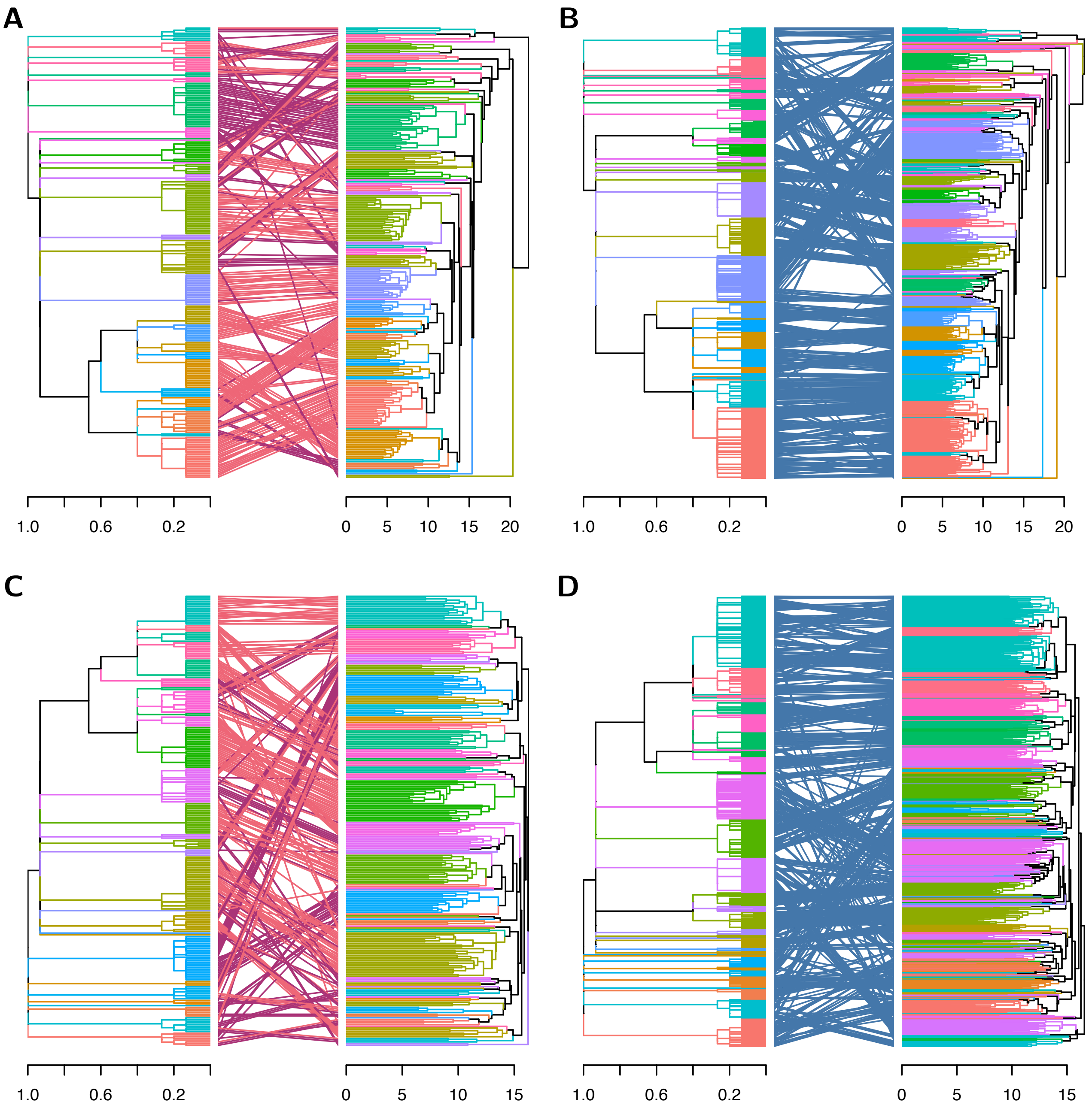


**S7 Fig. Concordance and discordance between virus genomic compositional features used by machine learning models and virus taxonomy.**

Tanglegrams compare hierarchical clustering of viruses by taxonomy (left in all panels) to clustering by (A–B) genome features or (C–D) model explanations from the combined genome feature-based model (SHAP values). Lines connect individual species in each dendrogram. Terminal branches are coloured by family, while connecting lines are coloured to indicate human-transmitted (purple), zoonotic (pink) and other viruses (blue). Note that the colours assigned to each family depend on the order of families in the dendrogram and have been optimized for distinguishability of neighbouring families, meaning colours do not match across panels.

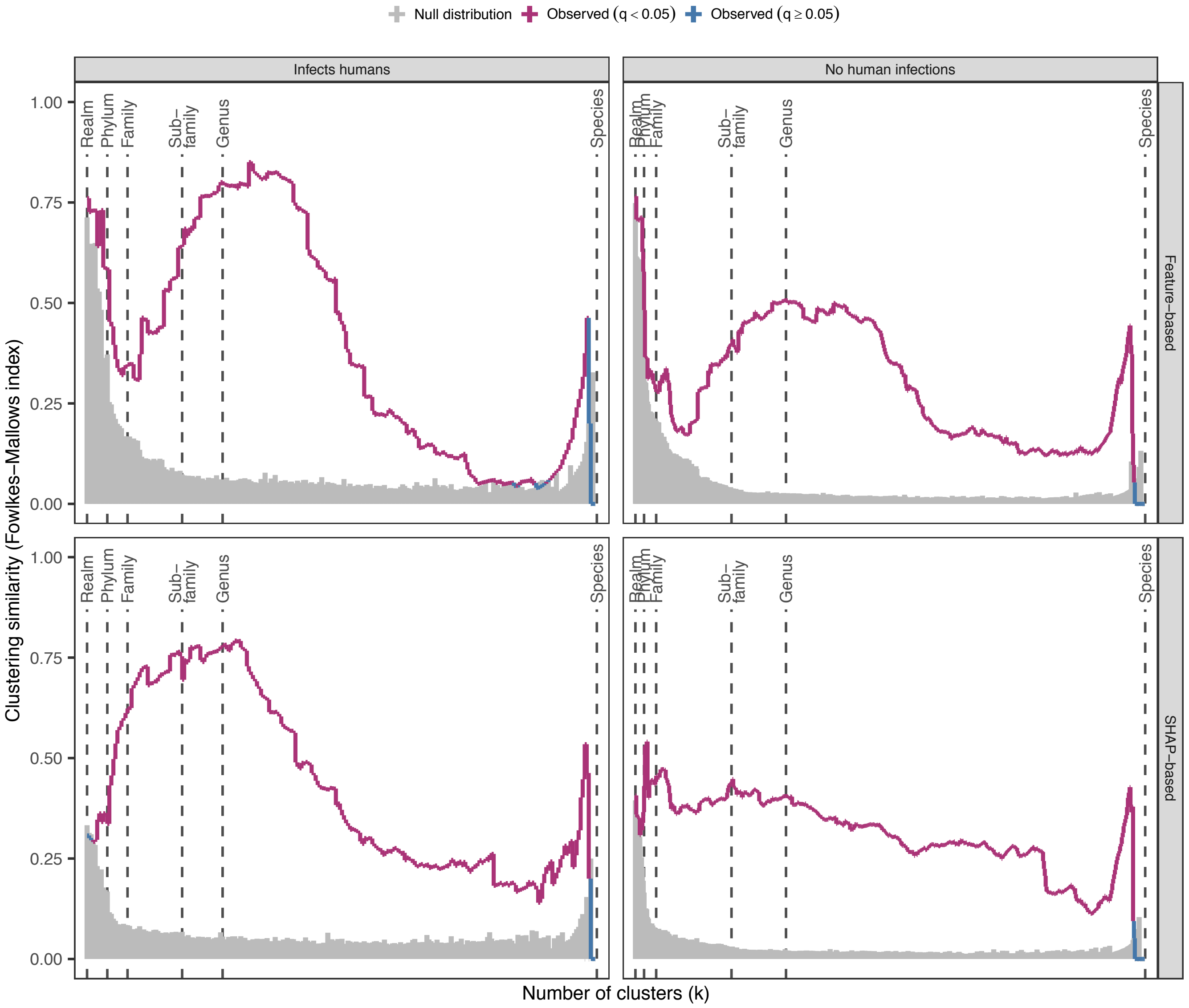


**S8 Fig. Association between virus-specific explanations of model predictions and taxonomy.**

Each panel measures clustering similarity (purple/blue line) when cutting virus taxonomy and either feature-based (top row) or SHAP value-based dendrograms (bottom row) into different numbers of clusters. Increasing the number of clusters (k) therefore compares clustering at increasingly shallower parts of the respective dendrograms (i.e. comparing more closely related viruses). A subset of taxonomic levels corresponding to different levels of subdivision are labelled for orientation. A clustering similarity of 1 would indicate complete agreement in the membership of all clusters, while a similarity of 0 indicates no agreement [11]. In all panels, empirical null distributions (grey) were obtained by randomly shuffling the labels of both dendrograms 1000 times. Clustering similarities significant at the 0.05 level (after Bonferroni correction) are illustrated in purple, while values which are not statistically distinguishable from the empirical null distribution are shown in blue. The correspondence between all dendrograms and virus taxonomy was generally higher than expected by chance. However, clustering of human-infecting viruses shows a lack of correspondence between how genome features are used in the model (as measured using SHAP values, bottom row) and the highest levels of virus taxonomy, even when such information was available amongst the input features (top row). Specifically, for both human infecting and non-human infecting viruses, correspondence between clustering and virus taxonomy declines from the Family to the Realm levels in SHAP-based clustering, but increases in feature based clustering. This indicates that while feature based clustering closely mirrors high level virus taxonomy, SHAP-based clustering links divergent viruses (cf. S7 Fig, panel C), implying similar feature usage even among viruses considered unrelated in the taxonomy.

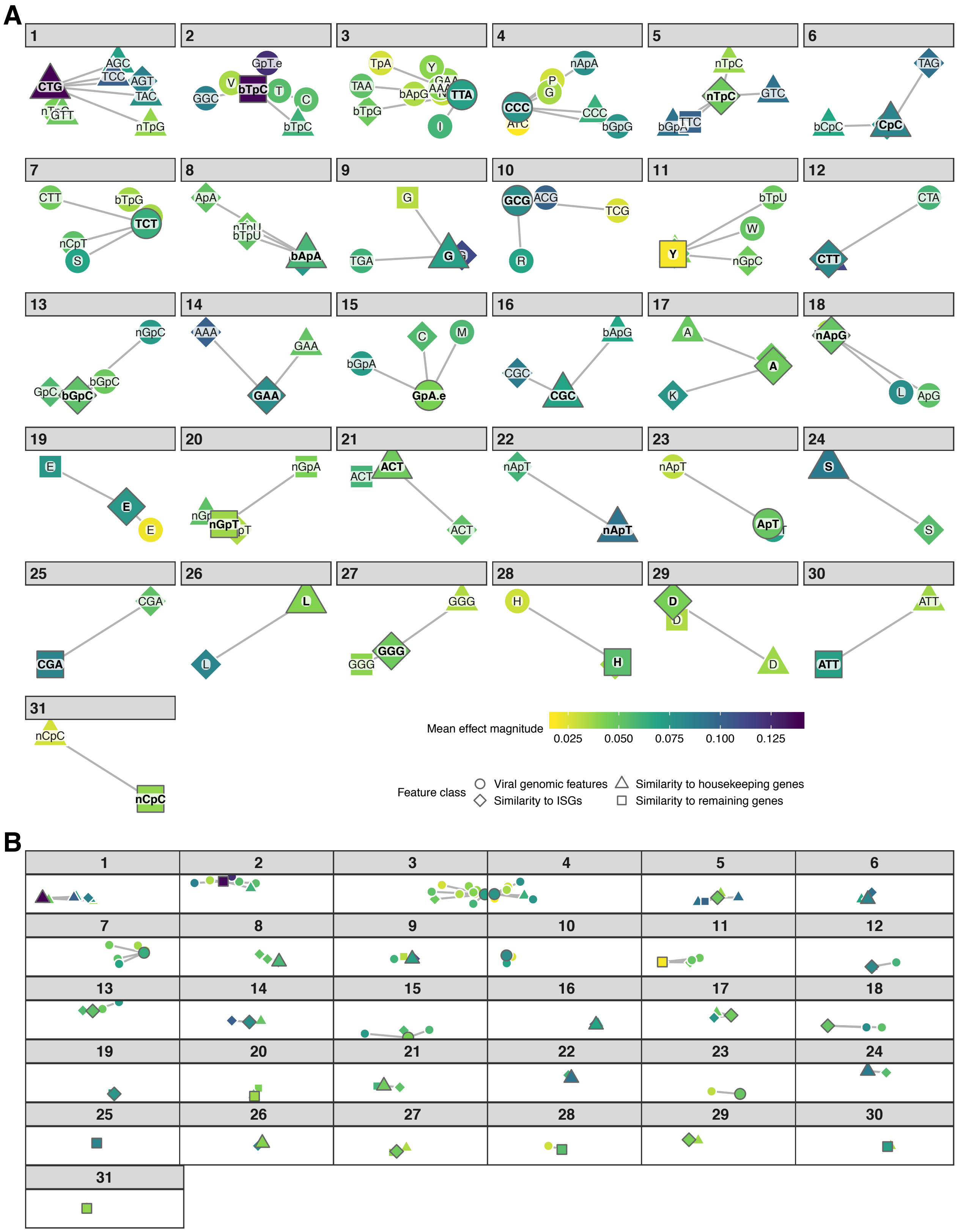


**S9 Fig. Clustering reveals high correlation between diverse feature types.**
Discrete clusters of features were obtained using affinity propagation clustering based on the Spearman correlation between all features present in the final model. Clusters are numbered to match their relative importance as defined in Fig 2D. Distances between features are illustrated in two dimensions, obtained by multidimensional scaling of the pairwise

correlation matrix. Individual clusters are shown to different scales in (A) for readability, while (B) shows all clusters on the same scale. All points are shown connected to the exemplar feature of that cluster, which is also indicated in bold font. Colours indicate the magnitude of each feature's effect on the combined genome feature-based model's output, calculated as the mean of absolute SHAP values across all viruses in the training data, and averaged across all 1000 model training iterations. Feature names abbreviated to a single letter indicate amino acid biases, while three-letter codes written in capital letters indicate codon biases. Dinucleotide biases are abbreviated in the form "CpG" and were calculated separately for codon bridge positions (abbreviation preceded by "b", e.g. "bCpG"), non-bridge positions (preceded by "n", e.g. "nCpG"), and also across all coding sequences of a given genome (no prefix, e.g. "CpG") or across the entire virus genome (suffix ".e", e.g. "CpG.e").

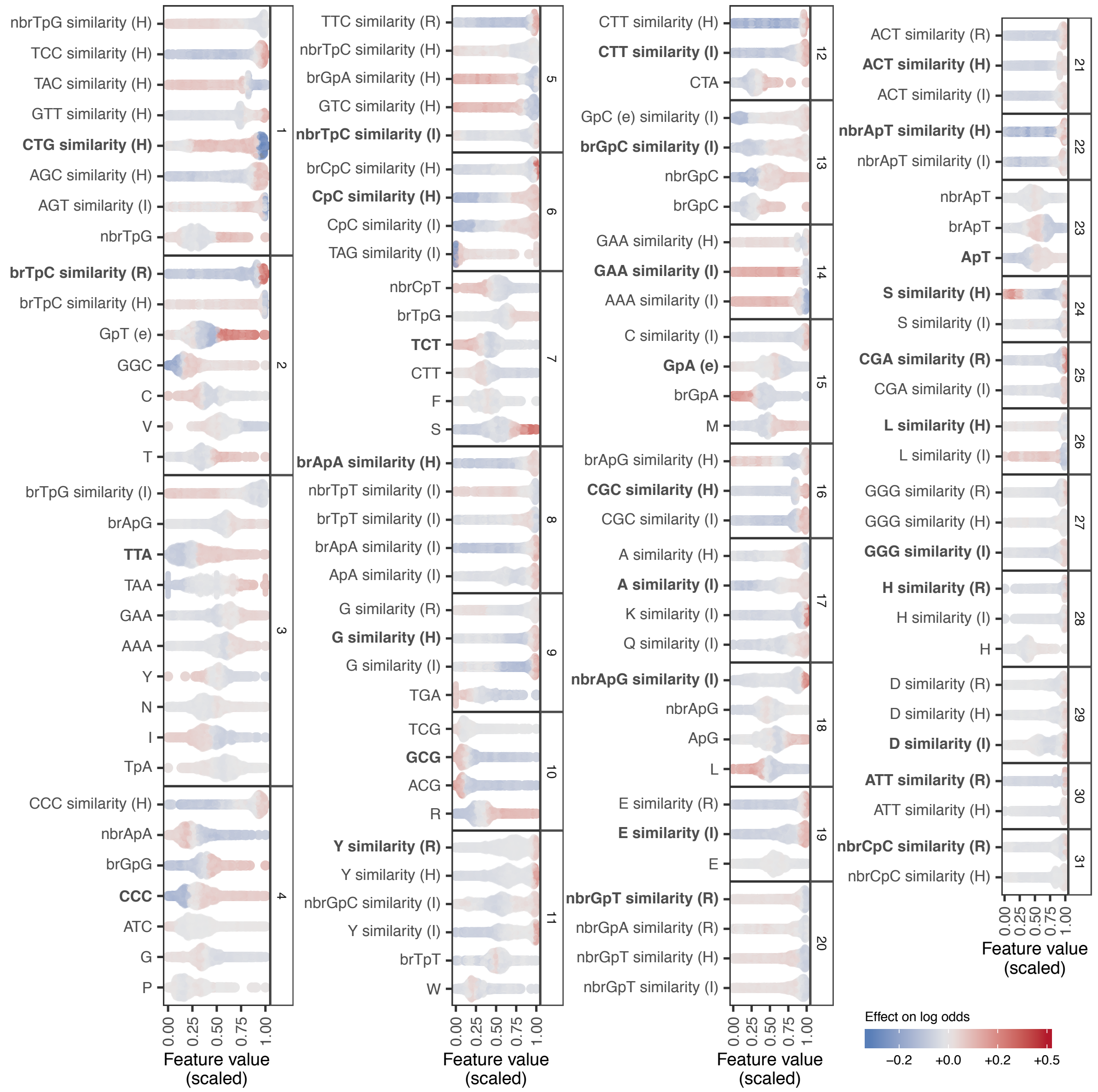


**S10 Fig. Directionality of relationships between genome composition feature values and the estimated odds of infecting humans.**

Each panel indicates a discrete cluster of correlated features, numbered by relative importance (see Fig 2D). Points are coloured to show the effect of each feature on the predicted log odds that an individual virus infects humans (i.e. the SHAP value for that feature for a given virus, derived from the combined genome feature-based model). The x-axis shows observed feature values (scaled to lie between 0 and 1 to place all features on the same scale), with points jittered to approximate local density where they overlap. Feature abbreviations follow a similar pattern to those in S9 Fig, except that bridge and non-bridge dinucleotide biases are preceded by "br" or "nbr", respectively. Capital letters in parentheses indicate the set of human genes used as baseline to calculate measures of compositional similarity (I = ISG, H = housekeeping genes, R = remaining genes). Names ending in "(e)" represent features calculated across the entire genome; all other features were calculated with reference to coding sequences only. The exemplar of each cluster is highlighted in bold.

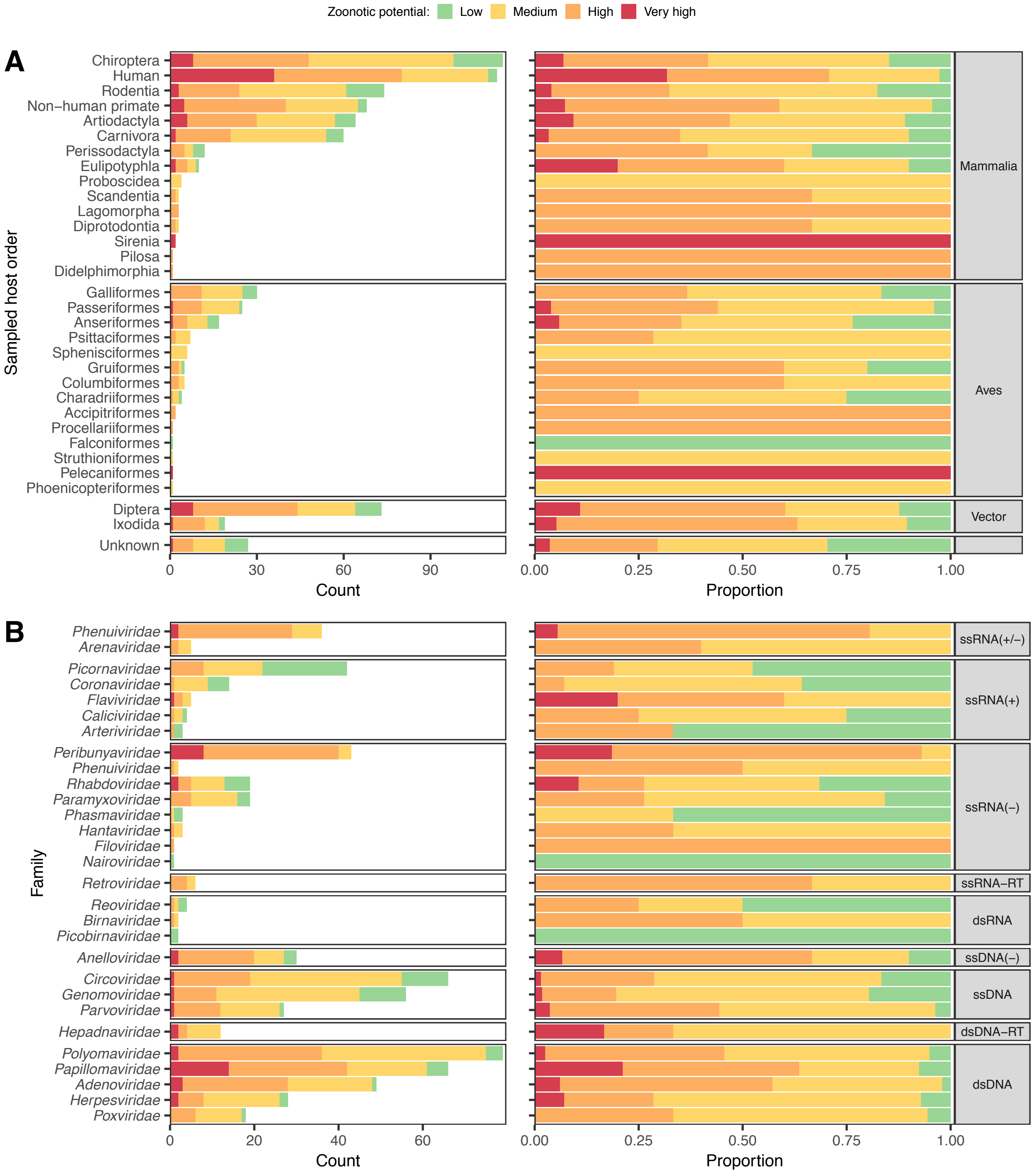


**S11 Fig. Distribution of zoonotic potential categories among 758 virus species which were not in the training data.**

(A) Distribution of zoonotic potential category assignments by sampled host, for all viruses in fig 3A. (B) Distribution of zoonotic potential categories among viruses from Fig 3A which were sampled from non-human hosts, arranged by viral family and genome type.

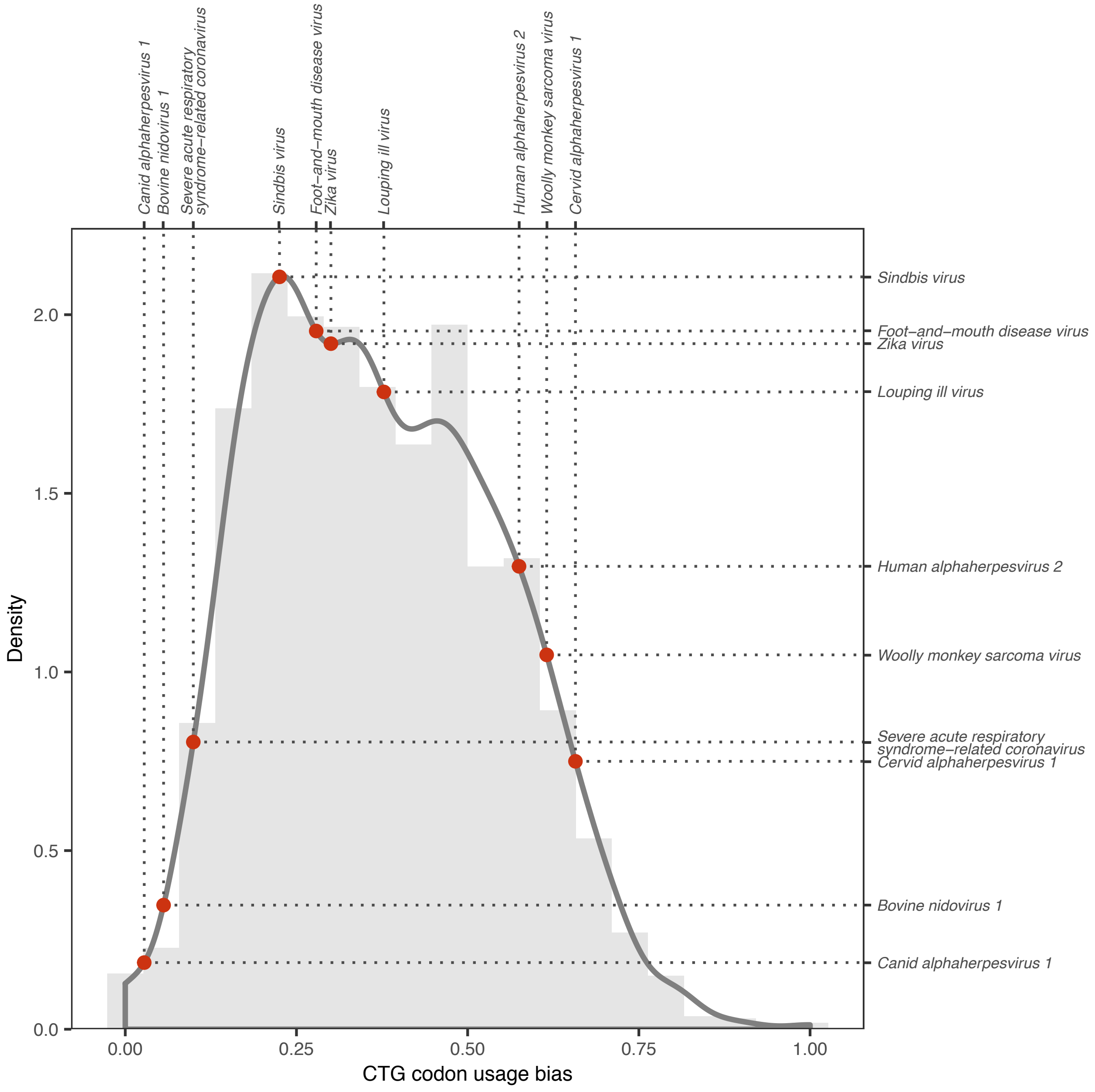


**S12 Fig. Illustrative calculation of genome feature similarity values.**

A histogram (light grey bars) shows the distribution of CTG codon usage bias observed among human non-ISG housekeeping genes. This distribution was used to estimate an empirical probability density function (dark grey line). Evaluating this function for the values of the same feature observed for specific viruses gave the final similarity scores (red dots, with the x-axis representing observed values and the y-axis the new similarity score). Using this similarity score as representing an estimate of how well each virus genome mimics the particular population of human genes resulted in re-arrangement of viruses which may help to predict the ability to infect humans. Because similarity scores were calculated via a density function, viruses with feature values significantly outside the range observed for a given set of human genes received a score of 0, but such cases were rare (0.025% of calculated similarity scores, affecting 5% of similarity-related features).

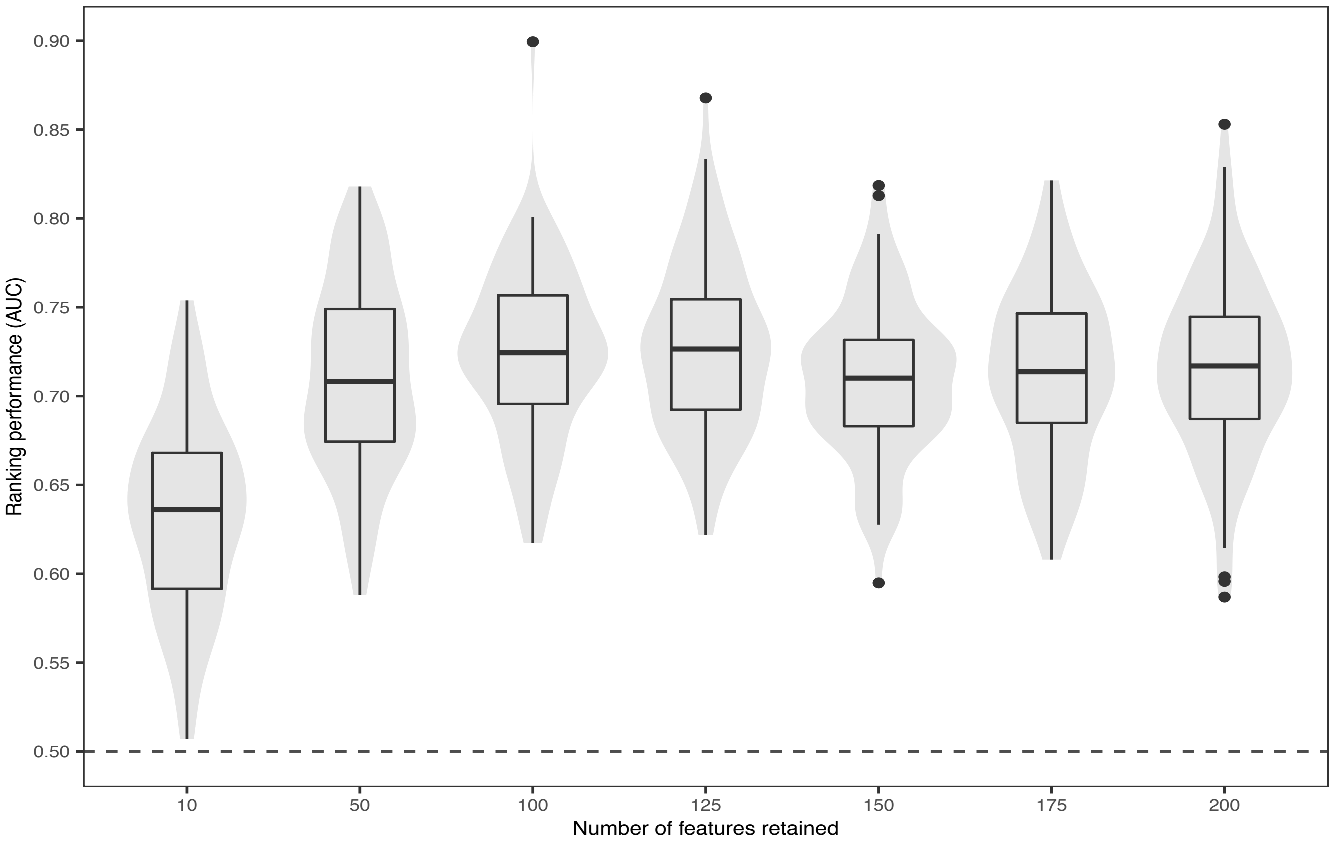


**S13 Fig. Ranking performance when training classifiers on restricted numbers of features, with all feature sets included.**

Boxplots and shaded areas illustrate the distribution of AUC values obtained when training classifiers on 100 random test:train:calibrate splits. For each set of classifiers, the top N most predictive features selected from amongst all feature sets were retained, with N indicated on the x-axis.

**S1 Table. (separate file)**

Predicted probabilities of human infection, zoonotic potential categories, and relative priority ranks for all viruses in the manuscript, derived from the combined genome feature-based model.
